# Tobacco smoke exposure is a driver of altered oxidative stress response and immunity in head and neck cancer

**DOI:** 10.1101/2024.10.17.618907

**Authors:** Y. Li, P. Yadollahi, F. Essien, V. Putluri, SRA. Chandra, K.R. Kami Reddy, AHM. Kamal, N. Putluri, LM. Abdurrahman, E. Ruiz-Echartea, K. Ernste, A. Trivedi, J. Vazquez-Perez, W.H. Hudson, W. Decker, R. Patel, A.A. Osman, F. Kheradmand, SY. Lai, JN. Myers, HD. Skinner, C. Coarfa, K. Lee, A. Jain, A. Malovannaya, M.J. Frederick, V.C. Sandulache

**Affiliations:** Bobby R. Alford Department of Otolaryngology Head and Neck Surgery, Baylor College of Medicine, Houston, TX; Advanced Technology Core, Dan Duncan Cancer Center, Baylor College of Medicine, Houston, TX; Dan L Duncan Comprehensive Cancer Center, Baylor College of Medicine, Houston, TX; Department of Molecular and Cellular Biology, Baylor College of Medicine, Houston, TX; Center for Precision Environmental Health, Baylor College of Medicine, Houston, TX; Department of Pathology & Immunology, Baylor College of Medicine, Houston, TX; Center for Cell Gene Therapy, Baylor College of Medicine, Houston TX; Department of Radiation Oncology, Baylor College of Medicine, Houston, TX; Department of Head and Neck Surgery, University of Texas MD Anderson Cancer Center, Houston, TX; Department of Medicine-Pulmonary Baylor College of Medicine, Houston, TX; Center for Translational Research on Inflammatory Diseases, Michael E. DeBakey Veterans Affairs Medical Center, Houston, TX; Department of Radiation Oncology, UPMC Hillman Cancer Center, Pittsburgh, PA; Verna and Marrs McLean Department of Biochemistry and Molecular Pharmacology, Baylor College of Medicine, Houston, TX; Mass Spectrometry Proteomics Core, Baylor College of Medicine, Houston, TX

**Keywords:** tobacco, Nrf2, oxidative stress, Keap1, glutathione, PDL1

## Abstract

**Purpose:** Exposomes are critical drivers of carcinogenesis. However, how they modulate tumor behavior remains unclear. Extensive clinical data link cigarette smoke as a key exposome that promotes aggressive tumors, higher rates of metastasis, reduced response to chemoradiotherapy, and suppressed anti-tumor immunity. We sought to determine whether smoke itself can modulate aggressive tumor behavior in head and neck squamous cell carcinoma (HNSCC) through reprogramming the cellular reductive state.

**Experimental design:** Using established human and murine HNSCC cell lines and syngeneic mouse models, we utilized conventional western blotting, steady state and flux metabolomics, RNA sequencing, quantitative proteomics and flow cytometry to analyze the impact of smoke exposure on HNSCC tumor biology.

**Results:** Cigarette smoke persistently activated Nrf2 target genes essential for maintenance of the cellular reductive state and survival under conditions of increased oxidative stress in HNSCC regardless of HPV status. In contrast to e-cigarette vapor, conventional cigarette smoke mobilizes cellular metabolism toward oxidative stress adaptation, resulting in development of cross-resistance to cisplatin. In parallel, smoke exposure modulates both expression of PDL1 and the secretory phenotype of HNSCC cells through activation of NF-κB resulting in an altered tumor immune microenvironment (TIME) in syngeneic mouse models and altered PBMC differentiation that includes downregulated expression of antigen presentation and costimulatory genes in myeloid cells.

**Conclusion:** Cigarette smoke exposome is a potent activator of the Nrf2 pathway and is a likely primary trigger for the tripartite phenotype of aggressive HNSCC consisting of: 1) reduced chemotherapy sensitivity, 2) enhanced metastatic potential and 3) suppressed anti-tumor immunity.

**Statement of significance:** The smoke exposome drives aggressive tumor behavior, treatment resistance and suppressed immunity through coordinated metabolic reprogramming. Successfully targeting this adaptation is critical to improving survival in smokers with head and neck cancer.

## Introduction

Head and neck cancer squamous cell carcinoma (HNSCC) is the sixth most prevalent cancer worldwide and is strongly linked to tobacco use, alcohol consumption, and human papillomavirus (HPV) infection (1). Patients with HPV-associated cancer generally experience excellent outcomes after radiotherapy or chemoradiotherapy compared to those with HPV-independent cancers (2). However, multiple prospective and retrospective analyses have shown that in cigarette smokers the survival advantage associated with HPV-associated oropharyngeal squamous cell carcinoma (OPSCC) is significantly reduced (3,4). Specifically, 5-year overall survival is 20-40% lower in smokers compared to non-smokers depending on HPV status (5,6).

Whereas most studies concerning tobacco smoke exposure have addressed the risk of carcinogenesis, our work has continuously focused on the impact of tobacco exposure, particularly continued at the time of cancer diagnosis and throughout treatment, on modulation of tumor biology, anti-tumor immunity and ultimately the effectiveness of chemoradiotherapy regimens (3), (4), (7). Tobacco smoke has now been linked to activation of indoleamine 2,3-dioxygenase 1 (IDO1) and immune modulation including increased infiltration of regulatory T-cells (Tregs) and reduced levels of cytotoxic T lymphocytes. (8) Furthermore, tobacco-associated carcinogens have been reported to directly induce chemoresistance through modulation of oxidative stress (9,10). Specifically, persistent exposure to tobacco can significantly heighten cellular oxidative stress, leading to continuous activation of antioxidant response element signaling (11) (12) (13).

We previously showed a link between tobacco smoke exposure to a distinct oncologic phenotype with higher rates of chemoradiotherapy failure in HNSCC, regardless of HPV association (3),(14). HNSCC patients with extensive tobacco smoke exposure, particularly those actively smoking at the time of diagnosis, demonstrate a suppressed, and potentially less functional anti-tumor immune response. (14) Using patient-level data, and preclinical HNSCC models, we subsequently linked hyperactivation of the Nrf2 signaling pathway with chemotherapy and radiation resistance in HNSCC (15), (16). This occurs largely through a metabolic adaptation to higher levels of chronic oxidative stress, resulting in an enhanced reductive state (higher levels of reducing equivalents) able to reset overall redox homeostasis. In the current study, we sought to determine whether Nrf2 signaling activated through tobacco smoke exposure could provide a common mechanistic link between altered immunity and reduced effectiveness of chemoradiotherapy through modulation of the oxidative stress response and metabolic adaptation in HNSCC regardless of HPV association.

## Methods

### Cell lines

A panel of human HPV-associated HNSCC (SCC090, SCC152, SCC 154, and UDSCC2) and HPV-independent cell lines (UMSCC22A, HN30 and HN31) was maintained in high glucose DMEM, supplemented with 10% fetal bovine serum, 1% glutamine, 1% vitamins, 1% non-essential amino acids, 1% penicillin/streptomycin, 1% sodium pyruvate, and 0.2% MycoZap™ Prophylactic (LT07-318, Lonza) to prevent mycoplasma overgrowth. Mouse MOC1 cells were cultured as previously described.(17)

Genetically engineered mouse models (GEMMs) of HPV-negative HNSCC were created using the Cre-LoxP system to induce tissue-specific alterations of genes, such as *TP53* and *PIK3CA* with or without *KEAP1*. These genes are among the most frequently altered in human HNSCC. Keap1-proficient (Keap1^+/+^) and Keap1-deficient (Keap1^fl/fl^) primary tumors isolated from GEMMs were digested using a Miltenyi tumor dissociation kit (Cat # 130-096-730), and a single-cell suspension was generated. *KEAP1*^+/+^ and *KEAP1*^fl/fl^ clones A, D, E, and F were provided by Dr. Rutulkumar Patel. *KEAP1*^+/+^ cell lines have intact copies of *KEAP1*, loss of both copies of *TP53* and harbor an activating *PIK3CA mutation (H1074R). KEAP1*^l/fl^ cell lines have one *KEAP1* copy deleted and the other floxed, as well as loss of both copies of *TP53* and activation of oncogenic *PIK3CA*. Clones A, D, E, and F were isolated from the *KEAP1*^l fl/fl^ cells. Cell lines were cultured at 37 °C in a humidified incubator with 5% CO_2_. Cisplatin resistant polyclonal populations utilized were previously described (18). Mycoplasma-free conditions were confirmed using the MycoAlert mycoplasma detection kit (Lonza, Basel, Switzerland). Cell line origin was confirmed by short tandem repeat (STR) analysis by the University of Texas MD Anderson Cancer Center Cytogenetics and Cell Authentication Core Facility every 3 months and for new cell line derivatives

Smoke-infused media was freshly prepared before each experiment by drawing smoke from 1 cigarette into 15 mL MEM medium over 1 minute, followed by sterile filtation. Cells were treated with the indicated concentration of smoke-infused media for the indicated times in acute smoke exposure settings. Chronic smoke exposure consisted of repeated exposure to increasing fractions of smoke-infused media from 0.5% to 4% depending on the cell line over a 6-month period, with application between 1-3 times per week.

Electronic cigarette vapor exposures were generated using 1.2 mL of liquid vape (nicotine concentration of 1.2 mg/mL, comparable to the nicotine content in one Marlboro Red cigarette) vaporized into 15 mL MEM medium.

### Western blotting

Cells were subjected to western blot analysis as previously reported (17) using the following antibodies: Nrf2 (PA5-27882, Invitrogen), GPX2 (MA5-36260, Invitrogen), KEAP1 (8407, Cell Signaling), NF-κB P65 (sc-8008, Santa Cruz Biotechnology), GPX2 (MA5-36260, Invitrogen), NQO-1 (ZRB1707, Millipore Sigma), β-actin (sc-81178, Santa Cruz Biotechnology). NE-PER Nuclear and Cytoplasmic Extraction Kit (78833, Invitrogen) was used to isolate the cytosol and nuclear portion according to the manufacturer’s instruction followed by the western blot analysis. Histone H3 (4499, Cell Signaling) was used as a nuclear fraction loading control, and α-tubulin (ab7291, Abcam) as the cytosolic fraction.

### Generation of *NFE2L2 (i.e, NRF2)* and *KEAP1* knockdown stable cell lines

GIPZ lentiviral human *NFE2L2* shRNA glycerol stocks (Clone Id: V2LHS_239104, V2LHS_64253), lentiviral human *KEAP1* shRNA glycerol stocks (Clone Id: V2LHS_254870, V3LHS_344984) and an empty vector (EV) control were obtained from Horizon Discovery. Lentiviruses were generated by transfecting HEK293T cells with shRNA or EV, envelope plasmid pMD2.G (#12259, Addgene) and the packaging vector psPAX2 (#12260, Addgene). Cells were infected with an EV or *NFE2L2* shRNA, *KEAP1* shRNA particles and selected for effective infection using 4 µg/mL puromycin; protein levels were confirmed by immunoblotting.

### Measurement of cellular redox states

Cells were labeled with 20 µM DCFDA (ab113851, Abcam) for 30 min and then treated with various concentrations of smoke-infused media for 1h. Cellular ROS levels were measured by fluorescence activated cell sorting (FACS) using BD FACS Aria flow cytometer.

### Cell cycle distribution analysis

HPV-HN30 and HPV+ UDSCC2 cells were seeded in 6 cm dishes and treated with various concentrations of smoke the following day for 24h and 48h. Cells were harvested and fixed overnight in 70% ethanol at 4°C. They were then stained with 20 µg/mL propidium iodide and 100 µg/mL RNase for 1h at 37°C, followed by analysis using a BD FACS Aria flow cytometer.

### FITC Annexin V apoptosis detection

Cells were seeded in 6 cm dishes and treated with various concentrations of smoke the following day for 48h. After treatment, cells were harvested and incubated in a buffer containing FITC Annexin V and propidium iodide (556547, BD Pharmingen™) for 15 minutes at room temperature (25°C) in the dark, followed by flow cytometric analysis.

### RNA-seq and ssGSEA analysis of TCGA cohort and cell lines

UMSCC47 and UDSCC2 cells were harvested after acute exposure to smoke-infused media for 8h, whereas SCC152 and SCC 154 cells were harvested following chronic smoke-infused media exposure with an additional bolus 8h prior to isolation. Total RNA from biological replicates was extracted using the RNeasy Mini Kit (74104, Qiagen) according to the manufacturer’s instructions and analyzed by whole transcriptomic sequencing (Psomagen Inc), which returned RSEM counts and effective gene sizes. Samples were normalized by calculating the FPKM-UQ values and then log transformed as log2 (X+0.01) for statistical analysis, except for single sample gene set enrichment analysis (ssGSEA) which was performed on data in linear space using the Broad Institute’s Gene Pattern website platform. Previously harmonized RNA-seq RSEM data from TCGA were downloaded from the Toil open-source data portal and normalized to derive FPKM-UQ values as we previously described (PMID: 36002187). For differential analysis of gene expression, only genes with an average log 2 FPKM value ≥ 2 for at least one group (i.e., treatment or control) were considered to avoid genes with low expression. Differentially expressed genes between cells treated with smoke and sham treated cells were analyzed with JMP 13 statistical software, which performs individual T-tests for each gene and applies a Benjamini-Hochberg correction (FDR<0.1) to calculate adjusted p-values. Fold-changes in gene expression refer to the ratio of geometric means (2^[log fold change]. Gene lists used for leukocyte and Nrf2 pathway ssGSEA score calculations were previously published and validated.(17) Essentially, cross-correlation coefficients of gene expression data were analyzed by two-way hierarchical clustering to identify modules of like expression, which were then validated by comparison with independent data or an orthogonal approach (17,19).

### Consensus hierarchical clustering

We previously described our in-house MATLAB script to perform consensus hierarchical clustering based on Ward’s linkage (17,19), which uses a modification of the resampling-based method published by Mont et al. Samples are clustered using Z scores derived independently for each gene (e.g., sample-wise Z score within a gene) and the already generated Z scores are transposed and re-used when clustering by gene. Our Matlab script is publicly available at https://github.com/aif33/Hierarchical-two-way-agglomerative-consensus-clustering.

### Proteomics

Cells were exposed to cigarette smoke-infused media for various periods and subjected to protein analysis. Protein extraction, digestion, and peptide fractionation were performed as previously described (20). Briefly, cells were lysed in 8M urea buffer, reduced or alkylated, and digested with LysC and Trypsin proteases. Twenty micrograms of peptide per sample was labeled with TMT16 pro plex isobaric label reagent (Thermo Fisher Scientific) according to the manufacturer’s protocol. High-pH offline fractionation was carried out to generate 24 peptide pools, which were acidified with a final concentration of 0.1% formic acid (FA). The deep-fractionated peptide samples were separated on a nano-LC 1200 system (Thermo Fisher Scientific, San Jose, CA) coupled to an Orbitrap Lumos ETD mass spectrometer (Thermo Fisher Scientific, San Jose, CA). The samples were loaded onto a 2 cm X 100µmI. D. Switched in-line with an in-housed 20 cm x 75µmI. D. column (Reprosil-Pur Basic C18, 1.9 µm, Dr. Maisch GmbH, Germany) equilibrated in 0.1% formic acid/water. The column temperature was maintained at 600C. Peptide elution was performed using a 110 min discontinuous gradient of 90% acetonitrile buffer (B) in 0.1%formic acid at 250nl/min (2-35%B, 86 min, 35-60%B, 6 min, 60-95%B, 9 min, 95-50%B, 9 min). The eluted peptides were directly electro-sprayed into a mass spectrometer operated in the data-dependent acquisition mode, acquiring HCD fragmentation spectra for a 2 s cycle time. MS1 was performed in Orbitrap (120000 resolution, scan range 375-1500m/z, 50ms Injection time), followed by MS2 in Orbitrap at 30000 resolution (HCD 38%) with the TurboTMT algorithm. Dynamic exclusion was set to 20 s, and the isolation width was set to 0.7m/z. Raw MS data processing, quantification, and differential analysis were performed as described previously (21). Reverse decoys and common contaminants were added to the NCBI human protein database (downloaded 2020_03_24) using Philosopher (22). Proteomics raw data files were converted to mzML using the MSConvert software. MASIC (23) was used to extract the precursor ion intensities for each peptide using the area under the elution curve as well as the reporter ion intensities. A smoothing method was used with a sampling frequency of 0.25 and an SCI tolerance of 10 ppm. Reporter ion tolerance was set to 0.003 Da, with reporter ion abundance correction enabled. Raw spectra were searched using MSFragger (v3.5) (24) and run with mass calibration. Settings included strict trypsin digestion for amino acids from 7 to 50 amino acids with up to two missed cleavages, clip protein N-term methionine, precursor ions with charge 2–6, precursor mass mode set to CORRECTED, and isotope error set to−1/0/1/2. For fragment spectra, TopN was set to 150, and the mass range was cleared between 125 and 135 to filter out reporter ions during spectral matching. The precursor mass error was set at ±20 ppm, and a fragment mass tolerance of ±0.02 Da. Static modification of C+57 carbamidomethylation and peptide N-terminus TMT16 label: +304.207. Dynamic modifications were set for methionine oxidation (+15.9949), acetylation (+42.0106), peptide N-terminal TMT16 label (+304.207), peptide N-terminal acetylation, and peptide N-terminal pyroGlu (−17.02650 for Q, C), and (−18.01060 for E). Multiple variable modifications were applied. Peptide validation was performed using a semi-supervised learning procedure in Percolator, as implemented by MokaPot (25). Peptides were grouped and quantified into gene product groups using gpGrouper (26). The resulting protein values were median-normalized and log-transformed. Heatmaps were generated using ComplexHeatmap (27). Data wrangling was performed with Python 3.8 (Python Software Foundation. Python Language Reference http://www.python.org), along with the third-party libraries Numpy31 and Pandas (mckinney-proc-scipy-2010, reback2020pandas). For proteomics data, group differences were assessed using the moderated t-test, as implemented in the R package limma (R version 4.2), and the false discovery rate was controlled using the Benjamini–Hochberg procedure.

Steady state and ^13^C flux metabolomics. Targeted steady-state metabolomic profiling was performed on cells exposed to smoke-infused media as previously described (16,28). To analyze the metabolic flux with [U-^13^C]-glucose, SCC1522 and UDSCC2 cells were plated in 10 cm dishes overnight. Subsequently, they were subjected to a 2h glucose starvation and incubated with media containing 12 mM [U-^13^C]-glucose in 0%, 5%, and 15% cigarette smoke-infused media for 72 h. Following incubation, the medium was aspirated and the cells were washed three times with cold PBS. An equal cell number of cells were harvested, snap-frozen in liquid nitrogen, and stored at -80 °C until metabolite extraction. [U-^13^C]-glucose-labeled cells were subjected to freeze-thaw cycles in liquid nitrogen, followed by homogenization in methanol: water (1:1) using needle sonication. The resulting samples were centrifuged for 10 min at 4°C (5000 rpm). Subsequently, the samples were filtered through a 3 K Amicon filter to eliminate proteins and lipids, dried using a speed vacuum, and reconstituted with a mixture of methanol and water (1:1). Glycolysis, TCA, and PPP intermediates and isotopomers were quantified using a Luna NH2 column (3 µm, 150 × 2 mm, Phenomenex, Torrance, CA) following a previously described method (28,29). For cysteine flux, the amino acid metabolites were separated through the XBridge Amide HPLC column (3.5 µm, 4.6 × 100 mm, Waters, Milford, MA) in both ESI positive (Method A). For ESI positive, mobile phases A and B were 0.1% formic acid in water and acetonitrile, respectively. Gradient flow: 0-3 min 85% B; 3-12 min 30% B, 12-15 min 2% B, 16 min 95% B, followed by re-equilibration till the end of the gradient 23 min to the initial starting condition of 85% B. Flow rate of the solvents used for the analysis is 0.3 mL/min. The injection volume was 10 µl.

For untargeted metabolomics, the cells were freeze-thawed and homogenized by needle sonication in 50/50 methanol/water. Three volumes of methanol/acetonitrile (50/50, v/v) were then added to the samples. The samples were vortexed for 5 min and kept at -20°C for 10 min. samples were then centrifuged for 10 min at 4°C and 15,000 rpm. The samples were then evaporated to dryness using a GeneVac EZ-2 Plus SpeedVac (SP Scientific). Aliquots were reconstituted in 100 µL of methanol/water (50/50, v/v). Twenty microliters (20 µL) of each supernatant were pooled and separated into aliquots for quality control (QC). The separation was performed using a Thermo Scientific Vanquish Horizon UHPLC system. A Waters ACQUITY HSS T3 column (1.8 μm, 2.1 mm x 150 mm) was used for reversed phase separation, and a Waters ACQUITY BEH amide column (1.7 μm, 2.1 mm x 150 mm) was used for HILIC separation. For the reverse phase, the gradient was from 99% mobile phase A (0.1% formic acid in water) to 95% mobile phase B (0.1% formic acid in methanol) for 21 min. For HILIC separation, the gradient was the same as that of solvent A: 0.1% formic acid, 10 mM ammonium formate, 90% acetonitrile, 10% H2O, and solvent B: 0.1% formic acid, 10 mM ammonium formate, 50% acetonitrile, and 50% H2O. For positive and negative modes of RP and HILIC, gradient flow: 0-3 min 1% B; 4-11 min 50% B, 12-21 min 95% B, followed by re-equilibration until the end of the gradient 30 min to the initial starting condition of 1% B. Both columns were run at 50 °C with a flow rate of 300 μL/min and an injection volume of 2 μL. A Thermo Scientific Orbitrap IQ-X Tribrid Mass Spectrometer was used for data collection with a spray voltage of 3500 V for the positive mode (reverse phase separation) and 2500 V for the negative mode (HILIC separation) using the H-ESI source. The vaporizer temperature and ion transfer tube were both 300°C. The capillary temperature and auxiliary gas heater temperature were set at 300 and 350°C, respectively. The sheath gas, auxiliary gas, and sweep gas flow rates were set to 40, 8, and 1 (in arbitrary units), respectively. For full MS, the data were acquired using an orbitrap detector with 120,000 resolution and quadrupole isolation. The scan range was set from 70 to 900 m/z, the normalized automatic gain control (AGC) target was 25%, and the absolute AGC value was 1.000e5 for positive and negative polarity. Compounds were fragmented by data-dependent MS/MS using HCD activation with HCD collision energies of 30, 50, and 150% via quadrupole isolation, and the detector type was orbitrap with 30,000 resolutions. The normalized automatic gain control (AGC) target was 100%, whereas the absolute AGC value was 5.000e5 for positive and negative polarity with a 50 ms maximum injection time. The acquired dataset was mined using Compound Discoverer (version 3.3.3.2, ThermoFisher Scientific, USA) for peak detection, integration, and identification. Metabolite identification was performed using the mzCloud and NIST 2020 high-resolution mass spectral (HRMS) library (Thermo Fisher Scientific), a retention time-based in-house mass spectral library containing 600 compounds, and the Human Metabolome Database (HMDB) mass list. A pooled quality control (QC) sample was run between every 10 samples during LC/MS analysis and used as a reference sample during metabolite identification using Compound Discoverer. For data analysis, the peak area was log2 transformed, and the median IQR method was normalized for each method. Differential metabolites were identified by performing a Student’s t-test to calculate p-values, followed by determination of the false discovery rate (FDR < 0.25) using the Benjamini-Hochberg method.

### Integrated metabolomic – transcriptomic analysis

Genes associated with differentially expressed metabolites were determined by using the Human Metabolome Database (HMDB)(30), as described previously (31,32). Enriched pathways based on differentially expressed metabolites were determined using over-representation analysis implemented in MSigDB (33) compared with the Hallmark and the Gene Ontology databases. A hypergeometric distribution test was used with significance defined as FDR-adjusted p-value<0.05. To integrate differentially expressed metabolites (DEMs) with differentially expressed genes (DEPs) from matching experiments, the DEGs were cross-referenced to genes associated with DEMs to create an overlapping signature.

### Hoechst assay and clonogenic assays

Hoechst and clonogenic assays were used to evaluate the cytotoxicity of cells exposed to smoke-infused media using established techniques.(15,16,18,34,35)

### ELISAs

Cells were seeded at the same density on day 1 and changed to fresh media the following day (day 2). The conditioned media were harvested after 24 and 48 h (days 3 and 4), and the total number of cells was counted accordingly. Secreted Prostaglandin E2 (PGE2, #514010) and Interleukin-6 (IL-6, #583371, #501030) levels were measured using commercial ELISA kits (Cayman Chemical, MI USA, R&D Systems, MN, USA) and normalized to the cell number. All data were repeated in triplicate and are represented as mean ± SD. P-values were calculated using Student’s t-test.

### Conditioned media experiments and flow cytometry

Two million parental HN30 cells and their corresponding 4% chronic smoke cells were seeded in 10 cm dishes followed by 4% smoke exposure or air bolus for 6 h the next day. Then, new media was substituted, and the supernatant collected after 24 h was sterile filteted to produce conditioned media. Human peripheral blood mononuclear cells (PBMC) were isolated from healthy donors, treated with conditioned media for 48 h, and then harvested for analysis by flow cytometry to characterize lymphoid and myeloid immune cell phenotypes. The myeloid cell flow panel consisted of CD11c-BV711 (BioLegend, San Diego, CA), CD68-PE (BioLegend), CD83-APC (BioLegend), HLA-DR/DP/DQ-FITC (BioLegend), CD1d-BV421 (BioLegend), CD80-BV605 (BD Biosciences, Franklin Lakes, NJ), and CD33-RB705 (BD Biosciences). The lymphoid cell panel consisted of CD3-APC-Cy7, CD4-BV605, CD8a-BV711, CD25-AF488, PD-1-PE, Lag3-BV421, and TIM3-APC (BioLegend). All flow events were acquired on an LSR II flow cytometer (BD Biosciences) and analyzed with FlowJo version 10.0.00003 for MacIntosh (Tree Star, Inc., Ashland, OR, USA).

### *In vivo* cigarette smoke exposure

All animal experiments were conducted in accordance with the U.S. Public Health Service Policy on the Human Care and Use of Laboratory Animals and the regulations of the Baylor College of Medicine Institutional Animal Care and Use Committee. To establish the chronic smoke *in vivo* model, six-to eight-week-old female C57BL/6J mice were exposed to an escalating dosage of cigarette exposure from 0.5 to 2.5 cigarettes per day for 8 weeks (exposure 5days/week, weekends off) accomplished by using a Airdyne2000 Air Compressor. In parallel, MOC1 cells were chronically exposed to smoke before orthotopic inoculation. Following the inoculation of 200,000 cells orthotopically (tongue) in each mouse, cigarette smoke exposure continued throughout the experimental period. Tumors were measured throughout the experiment and were harvested for transcriptional analysis. Murine PBMCs were isolated using a lymphocyte separation medium (#25-072-CV, Corning) for subsequent flow analysis (36).

### Statistical analysis

Student’s t-test and one-way analysis of variance (ANOVA) were used to assess the differences in means and variances in Excel. Statistical significance was set at p < 0.05. Additional statistical analyses are described, either above or within the individual experiments outlined below.

## Results

### Tobacco exposure is associated with altered immunity and Nrf2 hyperactivation in multiple smoking-associated malignancies including HNSCC

We previously published that overexpression of the Nrf2 downstream targets glutathione peroxidase 2 (*GPX2*) and aldo-ketoreductase family 1 (A*KR1C*) members were associated with a “cold” tumor immune microenvironment (TIME) in multiple smoking-related cancers (17) including OCSCC and non-small cell lung cancers (NSCLC). We also observed an increased frequency of *KEAP1/NFE2L2* pathway mutations in patients with cold tumors. Because *GPX2* has been described in the literature as the smoking-induced glutathione peroxidase (37), we decided to further investigate the relationship between smoke exposure, activation of the Nrf2 pathway and expression of its downstream targets, and immune phenotypes in HNSCC patient tumors and cell line models. To examine the correlation between smoking history and immune phenotypes, TCGA RNA-seq data from OCSCC patients was used to determine the distribution of tumors from individuals who smoke among patient clusters with increasing amounts of leukocyte infiltration. The ssGSEA scores from 14 different leukocyte subsets were used to cluster patients by immune profile (**Fig.1, Supplementary Table I)** using methods we previously published (17). Two adjacent clusters of patients with elevated presence of nearly every leukocyte subtype (e.g., “hot”) were easily discernable from the cold cluster with greatly reduced levels of all leukocytes. Tumors from patients who smoked concurrent with their diagnosis or had quit <15 years before were significantly enriched among the cold tumors (**Fig.1A** p<0.002). Next, we investigated the relationship between smoking history and Nrf2 pathway activation. We previously published a gene expression signature of Nrf2 activation comprised of 138 genes—many of which are known Nrf2 downstream targets —which was vetted through cross-correlation of gene expression from more than 9,000 TCGA tumor samples spanning dozens of different tumor types and validated by association with the presence of *KEAP1/NFE2L2* pathway mutations (17). Nrf2 activation scores in tumors derived from patients who recently quit smoking or were current smokers were significantly elevated compared to tumors from individuals who never smoked (p<0.0003 and p<0.02, respectively, **Fig.1B**). To dissect the specific contribution that mutations in the *KEAP1/NFE2L2* pathway had on this phenomenon, we repeated the analysis but stratified by mutation status (**Fig.1C**). Among tumors that were wild type for the *KEAP1/NFE2L2* pathway, Nrf2 activation (as measured by the gene expression signature) was still significantly elevated when patients had a recent or current smoking history compared to never smokers (p <0.02 and p<0.006, respectively). Although the percentage of tumors with *KEAP1/NFE2L2* pathway mutations didn’t vary much (e.g., 13%-18%) between never smokers (13%) and recent (18%) or current smokers (11%), there was a trend towards elevated Nrf2 pathway activation associated with mutations in current smokers that was significant (p<0.003) in recent smokers that suggests the mutations from the smokers may have had more functional impact. Collectively, the data from OCSCC patients suggests that smoking contributes to elevated Nrf2 activation in tumors through mechanisms both dependent and independent of pathway mutation.

**Figure 1.**
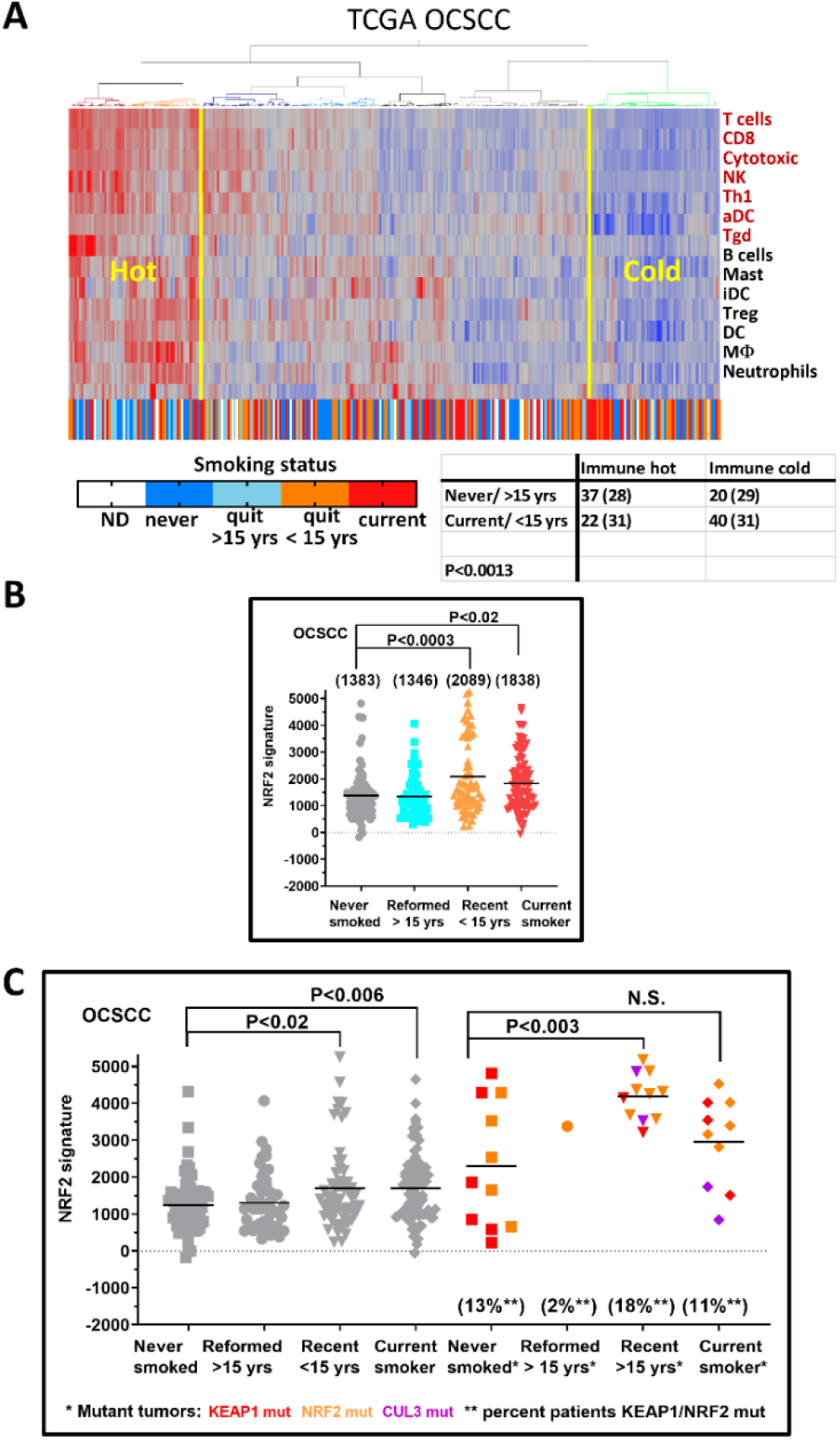
Smoking status is associated with TIME and Nrf2 activation in OCSCC patients. A) Heatmap of ssGSEA scores for 14 different leukocyte infiltrates after two-way hierarchical consensus clustering reveals enrichment of current or recently reformed (quit < 15 yrs before diagnosis) smokers among cold tumors (green right-most patient cluster). Smoking status of each patient is annotated across the bottom and the association between smoking status and immune phenotype was significant by Chi-square analysis (P<0.0013). B) RNA-seq data from the TCGA OCSCC cohort was used to calculate individual ssGSEA scores for each tumor sample using a 138 Nrf2 gene expression signature that is a measure of Nrf2 pathway activation. Patients were binned according to whether they never smoked, quit > 15 years before their cancer diagnosis (i.e., Reformed), quit <15 years before diagnosis (i.e., Recent) or still smoked (Current). C) Nrf2 scores for OCSCC patients were broken out separately for wild type tumor or those with *KEAP1/NFE2L2* mutations and according to smoking history. All p-values refer to comparisons to never smokers as the control group using a Dunnett multiple comparison test.

Outside the oral cavity (**Supp Fig.1A**), we also observed an increase in Nrf2 pathway activation in laryngeal/hypopharyngeal squamous cell carcinomas (LHSCCs) derived from current smokers that that nearly reached statistical significance in recently quit smokers compared to never smokers (e.g., average Nrf2 scores = 1642, 2004, and 1091). While NRF2 activation did not increase with tobacco use in HPV-associated oropharyngeal squamous carcinomas (OPSCCs), baseline levels of Nrf2 activation were elevated across HPV-associated OPSCC tumors, regardless of smoking status, compared to LHSCC and OCSCC tumors from patients who never smoked (**Fig.1; Supp Fig.1B**). Possibly, the presence of HPV may in and of itself lead to elevated Nrf2 activation. In LUAD (**Sup Fig.2A**), another smoking related cancer, Nrf2 scores were significantly elevated in tumors from individuals who recently quit smoking (p<0.05) or currently smoked (p<0.008) compared to never smokers. For this cancer cohort, the impact of *KEAP1/NFE2L2* pathway mutation appeared greater. Among *KEAP1/NFE2L2* wild type tumors (**Sup Fig.2B**), only those from current smokers had elevated Nrf2 scores compared to never smokers (p<0.05). While tumors with *KEAP1/NFE2L2* mutations maintained substantially elevated Nrf2 activity across smoking history categories (compared to wild type tumors), there was no relationship observed between Nrf2 activation levels and smoking history among mutants but the prevalence of *KEAP1/NFE2L2* pathway mutations was significantly reduced among never-smokers compared to individuals with a smoking history (**Sup Fig.2**).

### Tobacco smoke activates Nrf2 in both HPV-associated and HPV-independent HNSCC

To determine if the above patient level data denote merely a correlation, we sought out to mechanistically investigate the relationship between tobacco exposure and Nrf2 pathway activation. Acute tobacco exposure increased total cellular protein levels of Nrf2 in both HPV-associated (SCC090, SCC152, UDSCC2) and HPV-independent (UMSCC22A, HN30) human cell lines (**Fig.2A**). Upon re-exposure of the HPV-associated (UMSCC47) or HPV-independent (HN30) chronically smoke-exposed lines to acute smoke exposure, Nrf2 activation remained consistently elevated (**Fig.2B**). Acute (**Fig.3A**) and chronic (**Fig.3B**) exposure both resulted in localization of Nrf2 to the nucleus in a reactive oxygen species (ROS) mediated manner as demonstrated by reversal of the phenomenon when using the non-specific ROS scavenger N-acetyl cysteine (NAC) (**Fig.3C**). Nrf2 localization and its reversal by NAC correlated with relative *in vitro* cytotoxicity of cigarette smoke indicating that Nrf2 activation represents a survival response to exogenous ROS in HNSCC cells (**Fig.3D**) further supported by direct measurements of intra-cellular ROS levels following cigarette smoke exposure (**Sup Fig.3**).

**Figure 2.**
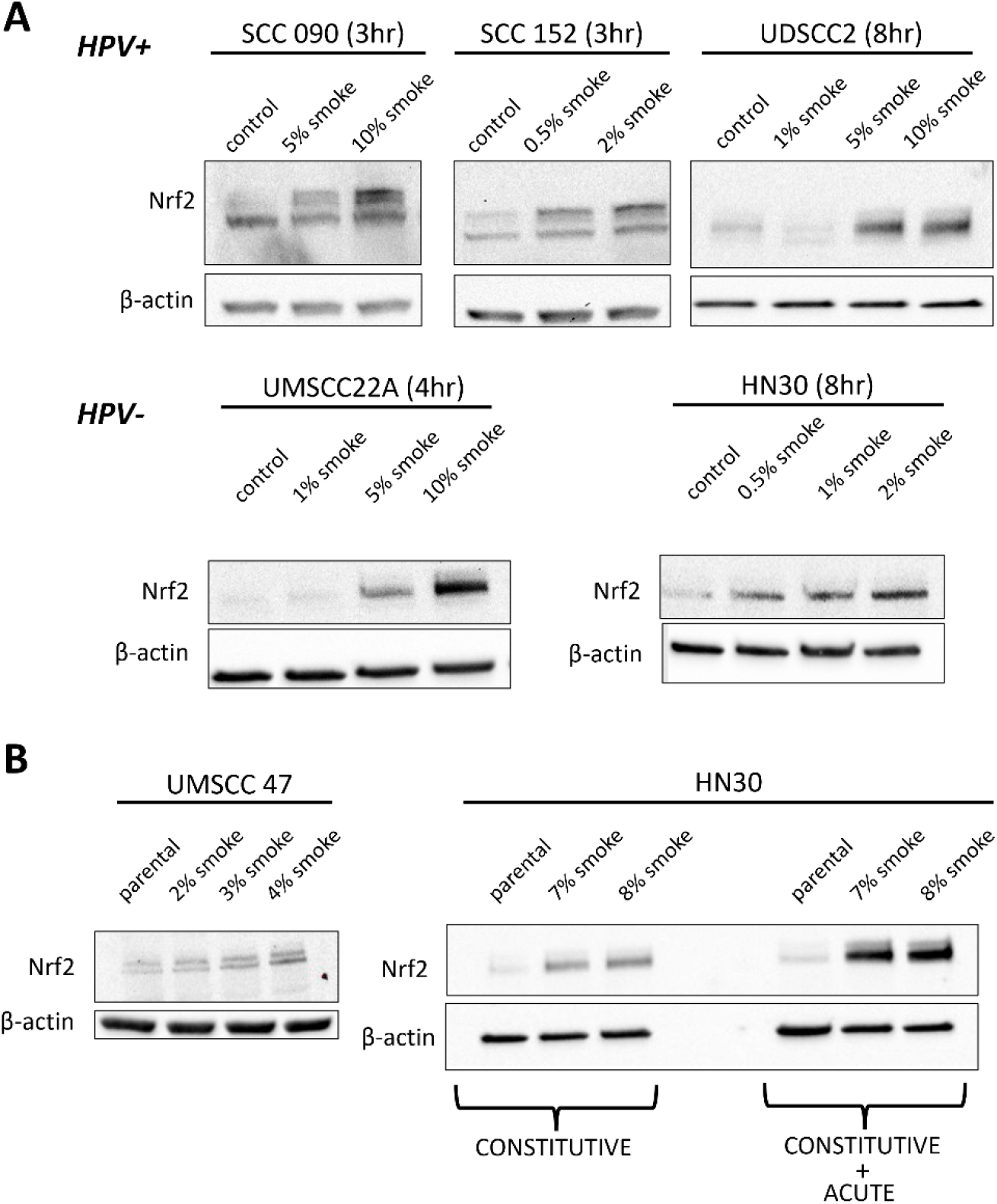
Tobacco exposure activates Nrf2 in both HPV-associated and HPV-independent HNSCC models. A) Nrf2 protein levels increased in a dose dependent manner upon acute smoke exposure at the indicated time periods in HPV-associated (upper panel) and HPV-independent (lower panel) HNSCC models. B) UMSCC47 and HN30 cells were chronically exposed to smoke-infused media at the indicated concentrations for 3-6 months; constitutive Nrf2 protein levels were measured. Right-panel lanes in HN30 cells indicate Nrf2 levels following an additional bolus of smoke 4h before harvesting. β-actin was used as the protein loading control.

To investigate the impact of smoke exposure on cell cycle progression, we employed flow cytometric analysis with PI staining to assess cell cycle distribution. Exposure to smoke caused increased G2/M phase arrest in HPV-independent HN30 cells after 24h and both S and G2/M arrest after 48h (**Supp Fig 4**) along with PARP and caspase-3 cleavage as well as increased Annexin V positivity. These changes coincided with evidence of increased Nuclear Factor kappa-light-chain-enhancer of activated B cells (NF-κB) activation which was further analyzed in additional detailed below. In contrast to conventional cigarette smoke, e-cigarette vapor exhibited lower cytotoxicity to HNSCC cells than conventional cigarette smoke even when nicotine levels were equivalent to those found in smoke-exposed HPV-associated (SCC152) and HPV-independent (HN31) cells (**Sup Fig.5**). To determine whether cigarette smoke exposure would activate Nrf2 and induce toxicity in normal mucosal tissue, we measured its effects in primary gingival keratinocytes which were found to be more susceptible to smoke exposure than HNSCC cells (**Sup Fig.6**).

We previously reported that evolution of cisplatin resistance was facilitated by Nrf2 pathway activation in three different HNSCC cell line genetic backgrounds (i.e., HN30, HN31, and PCI15B), which occurred both in the presence and absence of obvious *KEAP1/NFE2L2* pathway mutations and drove an enhanced ROS processing metabolic state (15). Consistent with an ROS-mediated mechanism of Nrf2 activation and cytotoxicity, we found that HNSCC cells adapted to higher levels of ROS through cisplatin exposure demonstrated profoundly lower cytotoxicity when exposed to cigarette smoke (**Sup Fig.7**). We then performed the converse experiment, by chronically exposing the HN30 and MOC1 parental cell lines to increasing concentrations of smoke-infused media, up to a maximum of 3% over a 9-month period. As shown in **SupFig.8**, chronic exposure to smoke-infused media significantly reduced the sensitivity of both cell lines to cisplatin. This could be attributed to the activation of Nrf2 levels following an additional smoke exposure in chronically exposed HNSCC cells, enhancing cell tolerance to ROS-induced cell death (**SupFig.9**).

### Nrf2-Keap1 homeostasis is relatively resistant to exogenous manipulation

Given the reasonably transient levels of Nrf2 activation following smoke exposure, we sought to exogenously manipulate Nrf2 in different cell backgrounds to further examine the relevant phenotype(s). Using 3 overlapping approaches, we demonstrated that although Nrf2 manipulation can impact phenotype, the manipulated steady-state is heavily selected against regardless of molecular methodology. As shown in **SupFig.10**, shRNA suppression of Nrf2 in our cisplatin resistant HN30R8 derivative which has high levels of Nrf2 and Nrf2 activation, using a commercial lentiviral construct results in substantial decrease in protein levels (whole cell and nuclear fraction) at early time points post infection, but the knockdown abated over time. Using multiple shRNA constructs (**SupFig.11**), we observed an inverse correlation between the re-establishment of Nrf2 expression in the cell lines and the suppression of cell proliferation in response to cisplatin treatment. Specifically, as Nrf2 expression levels increased, the inhibitory effect of cisplatin on cell proliferation was diminished, indicating a potential autoregulatory/compensatory of NRF2 mechanism at play.

Conversely, we used shRNA to suppress Keap1 levels in MOC1, a cell line with low basal Nrf2 protein expression, expecting a concomitant increase in Nrf2 levels. At early time points post infection, the concomitant increase in Nrf2 matched increased resistance to cisplatin, but both the Nrf2 levels and the phenotype were lost over time (**SupFig.12**). Similar results were observed in HN30 cells with shRNA targeting Keap1 at an early point. Nrf2 protein levels were significantly increased by the KD of Keap1 and exhibited significant protection against both smoke and cisplatin treatment (**SupFig.13A-C**). In contrast, we analyzed levels of Nrf2 and its downstream target Gpx2 in different clones of Keap1^fl/fl^ derived from GEMM tumors in which a single *KEAP1* copy was deleted and found stable and persistently high levels of Nrf2, Gpx2 and profound resistance to both smoke and cisplatin (**SupFig.14A-C**). Additionally, we overexpressed human *NFE2L2 (i.e*., *Nrf2)* in the HPV-independent HN30 cell line. Consistent with our previous findings, this overexpression resulted in the activation of Nrf2, demonstrated by its translocation into the nucleus and the upregulation of Nrf2-targeted gene expression (**SupFig. 15A and B**). Furthermore, Nrf2 OE conferred increased resistance to smoke and cisplatin treatments (**SupFig. 15C and D**). Similar results were observed in MOC1 cells overexpressing Nrf2 (**SupFig.16A-C**).

### Nrf2 activation by smoke generates an enhanced reductive state

To measure downstream cigarette smoke effects on Nrf2 target genes, we exposed two smoke-naïve HPV-associated HNSCC cell lines (UMSCC47 and UDSCC2) or two chronically smoke-exposed HPV-associated HNSCC cell lines (SCC152 and SCC154) to an acute bolus of 4% smoked media for 8h. In all four lines, smoke exposure dramatically increased Nrf2 activation scores (p <0.0001, **Fig.4**), consistent with our western blot results demonstrating increased levels of total cellular or nuclear Nrf2 protein (**Fig. 3**). Most Nrf2 signature genes (i.e.,75) were significantly overexpressed (**Supplementary Table II**) and only 24 (23%) were significantly decreased by at least 1.3-fold in one or more cell lines after smoke exposure (**Supplementary Table II**). There were 24 Nrf2 signature genes commonly increased by smoke ≥1.3-fold across all four cell lines (**Fig.4, Supplementary Table III**), with *AKR1C1, AKR1C3, GCLM, GGLC*, and *GPX2* among the top 10 genes with average fold change increases in expression from 3 to 9. Among the commonly upregulated 24 Nrf2 target genes, *AKR1C1, AKR1C3*, and *ALDH3A2* all function in lipid peroxide detoxification while *GPX2* reduces hydrogen peroxides as well as fatty acid hydroperoxides. Also elevated were *SLC7A11, GLCM, GCLC*, and *CHAC1* which are all functionally linked to maintaining glutathione levels and the antioxidant *SRXN1* (**Supplementary Table III**). Suppressors of ferroptosis (*STC2*) and NF-κB levels (*SQSTM1*) were among Nrf2 signature genes elevated following smoke exposure (**Supplementary Table III**) along with the autophagy-related gene *OSGIN1* (**Supplementary Table III**). A total of 11 Nrf2 targets were significantly elevated in cisplatin resistant cells from all three genetic backgrounds and in all four HPV-associated HNSCC cell lines stimulated by smoke (**SupFig.17**). This robust set of elevated Nrf2 target genes (**SupFig.17; Supplementary Table III)** included genes functioning in lipid peroxide detoxification (*AKR1C1, AKR1C3, GPX2*), glutathione and redox homeostasis (*GCLC, SLC7A11, TXNRD1*), autophagy (*OSGIN1*), *WNT* signaling (*FZD7*), and xenobiotic detoxification (*UGT1A6*).

**Figure 3.**
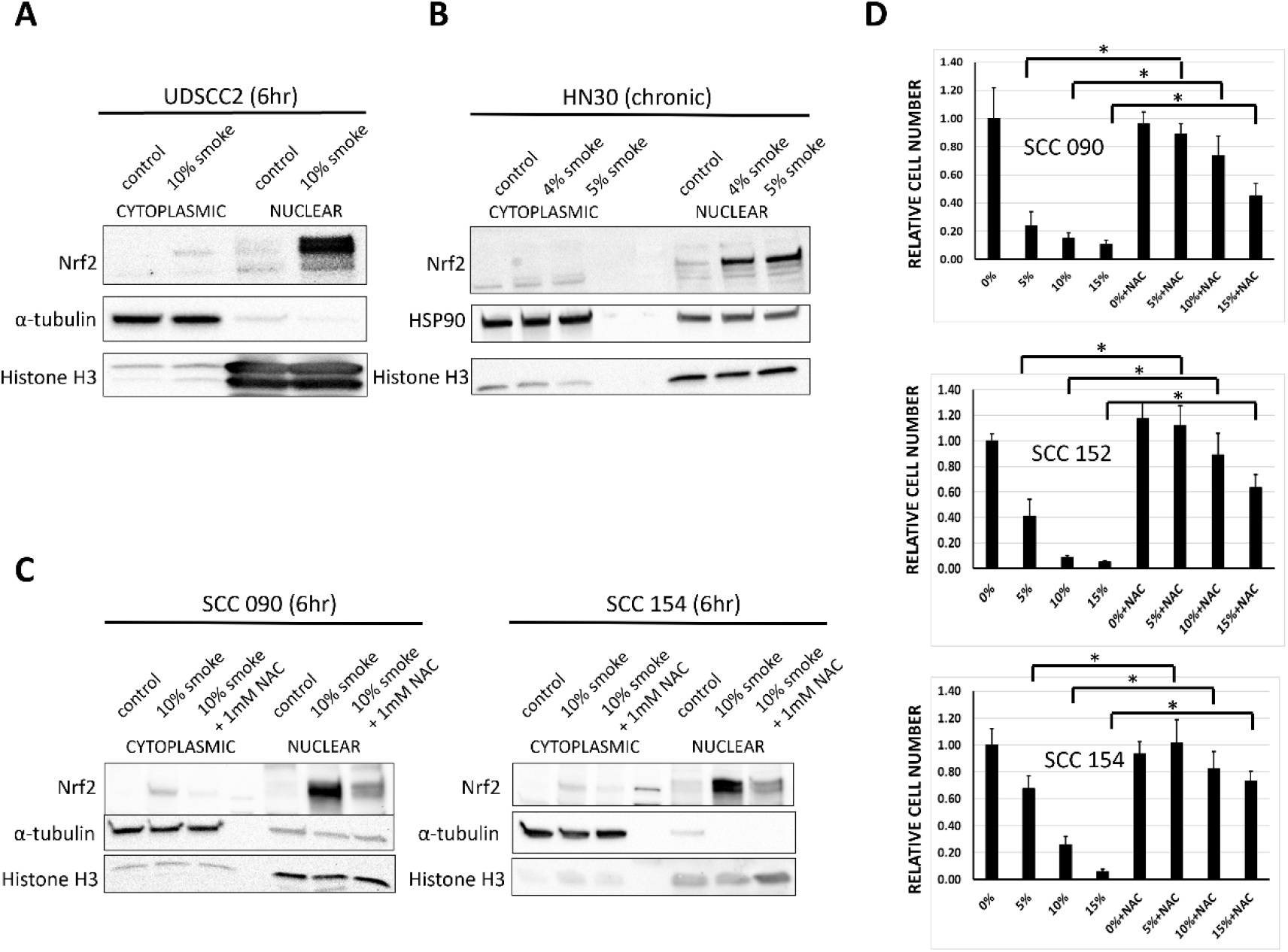
Tobacco exposure effects are ROS-mediated. A, B) Nrf2 protein levels were measured in nuclear and cytoplasm fractions after acute (6h) smoke exposure in UDSCC2 cells and in the chronic exposure model of HN30 cells. α-tubulin or HSP90 served as loading controls for the cytoplasmic fraction, and histone H3 for served as loading control for the nuclear fraction. C) Pretreatment with the ROS scavenger NAC inhibited smoke-induced Nrf2 nuclear translocation in acute smoke exposure models of SCC090 and SCC154 cells. D) SCC090, SCC152, and SCC154 cells were exposed to 0%, 5%, 10%, and 15% cigarette smoke-infused media in the presence or absence of 3mM NAC. Relative cell number at the end of a 72h exposure period was assessed using Hoechst assay. * denotes p-value <0.05 for the compared values.

**Figure 4.**
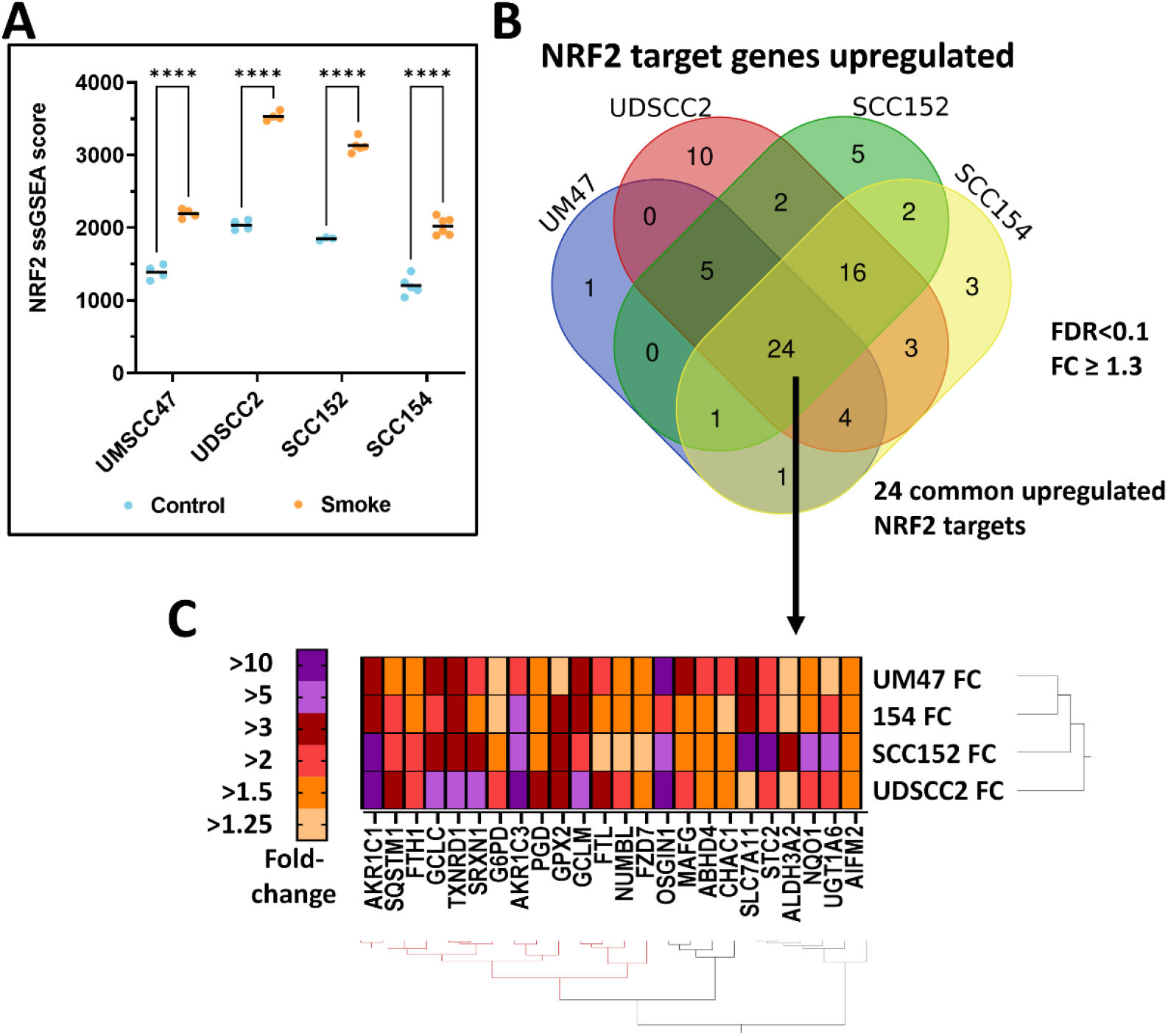
The Nrf2 pathway is activated in HPV-associated HNSCC cell lines exposed to smoke. A) NRF2 activation scores are significantly higher in both smoke-naïve (UMSCC47 and UDSCC2) and chronically exposed cell lines (SCC152 and SCC154) following an 8 hr bolus of acute smoke. **** adj P <0.0001. B) Venn diagram illustrating the number of overlapping Nrf2 signature genes (i.e., downstream targets) significantly elevated by at least 1.3 fold (e.g., FDR <0.1) after smoke exposure in the 4 HPV+ cell lines. C) Heatmap of fold change increases for the 24 commonly upregulated genes following two way hierarchical clustering, with color gradient reflecting the range of fold increase.

We identified 46 genes that were not downstream of Nrf2 but were upregulated by ≥ 1.3-fold in all 4 cell lines treated with smoke and 216 upregulated in at least 3 cell lines exposed to smoke (**Supplementary Table IV**). Genes upregulated in all cell lines were significantly enriched for regulation of cell proliferation, protein metabolism, apoptosis and response to stress (**Supplementary Table V**). Among the genes commonly upregulated lines we identified: (1) *CDKN1A*/p21 which mediates cell growth arrest; (2) heat shock proteins *HSPA1A* and *HSPA1B* which mediate cell survival to stress; and (3) PIM1 and PIM3, two pro-survival kinases that inhibit apoptosis. A total of 76 genes were significantly decreased by ≥ 1.3-fold in all HPV-associated cell lines after smoke treatment (**Supplementary Table IV and VI**) and these were enriched for genes regulating nucleosome organization, mucosal immune response, and innate immune response in mucosa (**Supplementary Table V**)-all of which were driven by downregulation of various histone genes. Among these were *H2BC10*,11,12, and 12L which have been directly linked with innate immunity (38). In all, there were roughly 40 different histone proteins downregulated in all lines following smoke treatment (**Supplementary Table VI**), although *pathways* tied to proliferation were noticeably absent (**Supplementary Table VI**).

To validate the transcriptomic data, we performed mass spectroscopy analysis of UDSCC2 cells acutely exposed to either 5% or 10% cigarette smoke-infused media and confirmed that the Nrf2 signaling pathway was most profoundly enriched. This was followed by the P53DN.V1 pathway consistent with the expected inactive functional status of p53 in an HPV-associated HNSCC cell line (**SupFig.18; Supplemental Table VII**). A similar analysis with SCC152 chronically exposed to smoke and restimulated with an acute bolus of smoke identified the Nrf2 pathway among the most upregulated. These cells also showed upregulation of the G2M checkpoint (Hallmark pathway)―consistent with a shift from rapid proliferation toward enhanced biomass generation―allowing for survival under stress conditions as we previously demonstrated in the context of cisplatin (**SupFig.19; Supplemental Table VIII**).

### Metabolomic shifts following smoke exposure support an enhanced reductive state

Steady state metabolomic analysis of UDSCC2 (**Supplemental Table IX**), SCC152 (**Supplemental Table X)** and SCC154 (**Supplemental Table XI**) demonstrated a shift in metabolite levels toward an altered reductive state inclusive of higher levels of NAD, homocysteine, glutathione, and increases in the 3-carbon glycolytic intermediates glycerol-3-phosphate and phosphoenolpyruvate. These changes echoed the transient and permanent alterations in the reductive state we previously linked to cisplatin induced oxidative stress (**SupFig.20**). Integration of metabolomic and transcriptomic data was performed to hone in on those metabolic enzymes most consistently and profoundly impacted by exposure to tobacco in HNSCC. As shown in **SupFig.21** and summarized in **Supplemental Table XII**-UDSCC (FDR 0.05), **Supplemental Table XIII-**SCC152 (FDR 0.05), **Supplemental Table XIV**-SCC154 (FDR 0.05), the greatest impact of tobacco exposure occurred in pathways specifically designed to restore the reductive state through recycling of glutathione and other primary and secondary reducing equivalents. Of note, the 2 enzymes consistently identified across all tested backgrounds were Gpx2 and Gsr (glutathione-disulfide reductase).

In SCC152 cells, smoke increased flux of carbon from ^13^C isotopically labeled glucose towards the PPP pathway, evident in elevated levels of labeled ribose/ribulose/xylulose-5-phosphate and sedoheptulose-7-phosphate. UDSCC2 cells exhibited a shift in glucose metabolism towards both the TCA and PPP pathways (**SupFig.22**). In contrast to carbon from glucose, carbon from ^13^C-cysteine was not differentially shunted under conditions of smoke exposure (data not shown). To further expand our understanding of these metabolic shifts we performed for the first time, unbiased metabolomics (**Fig.5**) of HNSCC cells exposed to acute cigarette smoke and e-cigarette vapor. As summarized in **Supplemental Tables A & B** we detected a significant shift in metabolites reflective of the overall reductive state inclusive of reduced glutathione. Although unbiased metabolomic analyses can detect extensive portions of the non-biologic exposome that are not readily analyzable, one component of the exposome of critical relevance to the current experiment was nicotine which was readily detectable using this approach and confirmed that e-cigarette vapor exposure resulted in clearly detectable nicotine levels despite a substantially reduced impact on the overall level of oxidative stress.

**Figure 5.**
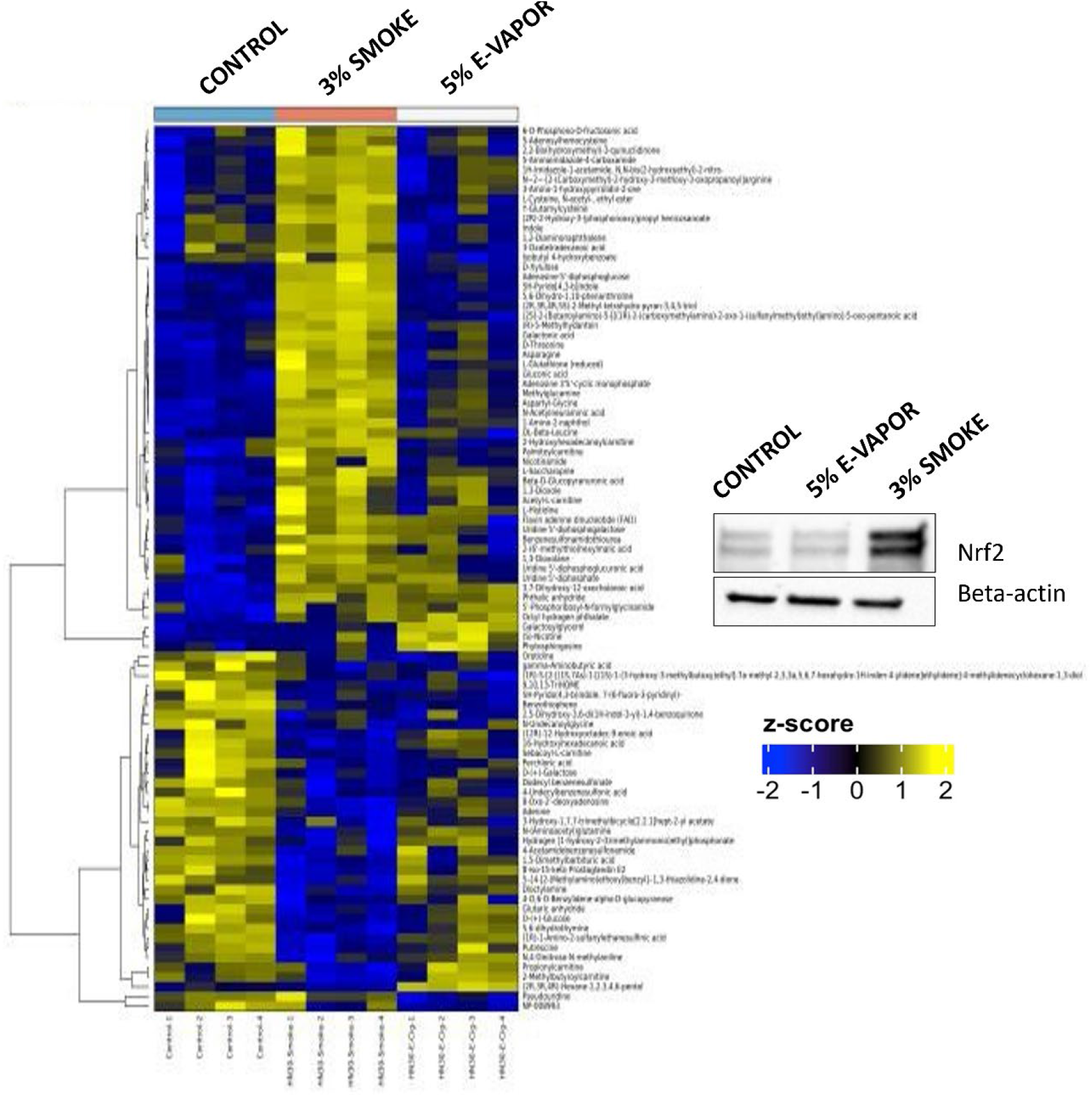
Acute smoke but not e-cigarette vapor exposure activates metabolic shifts neutralizing oxidative stress. Acute smoke exposure (8hr) of HN30 resulted in metabolic shifts, focused on pathways designed to neutralize oxidative stress and rebuild reducing equivalents. Data shown for metabolites with a differential FDR < 0.25 using unbiased metabolomics approach. In contrast, e-smoke exposure had a substantially diminished metabolic impact consistent with reduced oxidative stress. Inset demonstrates total cellular Nrf2 and β-actin protein levels measured at the time of harvest for metabolomic analysis.

### Nrf2 activation and immune modulation

Given the shifts in gene expression following Nrf2 activation by smoke exposure, we sought to measure its impact on inflammatory mediator production and immune cell function, *in vitro, in vivo* and using conditioned media approaches. As shown in **SupFig. 23**, a bolus of smoke exposure transiently activated Nrf2 expression in chronic smoke-exposed HN30 and UDSCC2 cells. However, PDL1 levels were substantially increased in both cell lines even after Nrf2 protein levels returned to baseline. We next investigated whether Nrf2-Keap1 homeostasis is coordinated with NF-κB, which we previously linked to altered *GPX2* activation (17). In the absence of smoke bolus, ectopic expression of Nrf2 led to increased expression of downstream NQ01 and paradoxically increased NF-κB in both HN30 (**SupFig. 15A**) and MOC1 (**SuppFig. 16A**), including nuclear accumulation in the former (**SupFig. 15B**), which can be attributed to reductive stress or reactive nitrogen species following excessive antioxidant activity. This was accompanied by significant increase in prostaglandin E2 (PGE2) after Nrf2 overexpression in both HN30 and MOC1 cells (**SupFig. 15E and 16C**). Elevated Nrf2 and PGE2 was also mirrored in the KEAP1 knockout GEMM cell lines (SupFig 14A and D). In chronically smoke-exposed HN30 cells (**SupFig. 24**), the accumulation of NF-κB in the nucleus corelated with increased production of PGE2 and IL-6. In contrast, acute smoke exposure led to elevated Nrf2 with a reduction in PGE2 levels for both HN30 and UDSCC2, that were consistent with failure of NF-κB translocation to the nucleus (**SupFig. 25A-B, D-E)**. However, no significant changes in IL-6 were found with the acute smoke exposures. Collectively, the data show that nuclear accumulation of NF-κB correlated is likely driven by chronic cellular stress and correlated with increased inflammatory mediators PGE2 and IL-6; whereas Nrf2 likely counters this effect in a more acute setting.

We performed duplicate *in vivo* experiments to recapitulate chronic smoke exposure using the murine oral cancer cell line MOC1 (39,40), which was implanted orthotopically in syngeneic immunocompetent mice. MOC1 cells, exposed to cigarette smoke (final concentration, 4%) for a period of >4 months were inoculated into mice that had also been chronically exposed to cigarette smoke and the smoke exposure was continued *in vivo* until completion of the experiment. Control MOC1 cells, never exposed to smoke were inoculated in mice with no cigarette smoke exposure. Variable rates of tumor growth were noted, with no clear change in tumor growth velocity attributable to smoke exposure. To determine whether the chronic smoke exposure altered the intrinsic biology of the tumors or the tumor immune microenvironment, RNAseq was used to compare 8 control tumors and 12 smoke exposed tumors. As shown in **SupFig. 26** and **Supplemental Table XV**, significant shifts in immune related genes were noted in smoke exposed tumors including decreased levels of chemokines *Cxcl5* (neutrophil recruitment) and *Ccl25* (T cell recruitment). Conversely, genes up-regulated in smoke exposed tumors mapped to GO pathways related to: 1) negative T cell selection, 2) regulation of NK cell mediated immunity and 3) negative regulation of lymphocyte mediated immunity among others. Although upregulation of Nrf2 itself was not detectable under these in vivo conditions, there was significant up-regulation of Nrf3 and the downstream target *Akr1C18* which has been previously linked to reduced chemotherapy response in preclinical murine models.

Given the difficulty of capturing the complete human smoking phenotype using a murine model, we used a simpler conditioned media exposure model to ascertain the effects of smoke exposure on the interaction between HNSCC cells and PBMCs. Two different approaches were utilized. First, bulk RNAseq was used to examine changes in the transcriptional signature of PBMCs following incubation with conditioned media that had been collected from HNSCC cells previously treated with a bolus of smoke followed by replacement with fresh media. As summarized in **Supplemental Table XVI**, genes significantly down-regulated in PBMCs incubated with smoke exposed conditioned media (i.e., SECM) from tumors mapped to GO pathways inclusive of: 1) antigen processing and presentation, 2) MHC protein complex assembly and 3) positive regulation of immune effector process among others. In a second approach, flow cytometry was instead used to characterize shifts in PBMC fate triggered by incubation with SECM. SECM treatment led to significant shifts in PBMC differentiation as shown in **SupFig.27** and **Supplemental Table XVII**, which included dramatically lower expression of MHC class II only within the myeloid CD11c^+^ (and not CD19^+^, not shown) antigen presenting cell population (***p<0.001), substantially lower CD80 (costimulatory) and CD1d (lipid antigen presentation) expression levels (***p<0.005).

## Discussion

Oxidative stress management plays a crucial role in both tumor formation and the effectiveness of anticancer treatments (41). Historically, Nrf2 was thought to primarily function as a tumor suppressor by regulating antioxidant gene expression and detoxifying reactive oxygen species (ROS) (42). However, recent studies have challenged this perspective using new evidence. ROS and the metabolic adaptations which accompany ROS exposure, have now been shown to mediate cellular reprogramming by selecting for aggressive features, such as enhanced invasiveness and resistance to therapy. We recently linked hyperactivation of Nrf2 through both mutational and transcriptional mechanisms to acquisition and maintenance of resistance to chemotherapy and radiation, the two mainstay treatments for HNSCC as well as for many other solid tumors (**Fig. 6**) (15), (43). Whereas selection of inactivating *KEAP1* or activating *NFE2L2* mutations in the context of treatment-induced oxidative stress can be detected in both preclinical models and human tumors, there remains an important question, namely whether some tumors are predisposed to hyperactivation of this pathway and development of subsequent treatment resistance.

**Figure 6.**
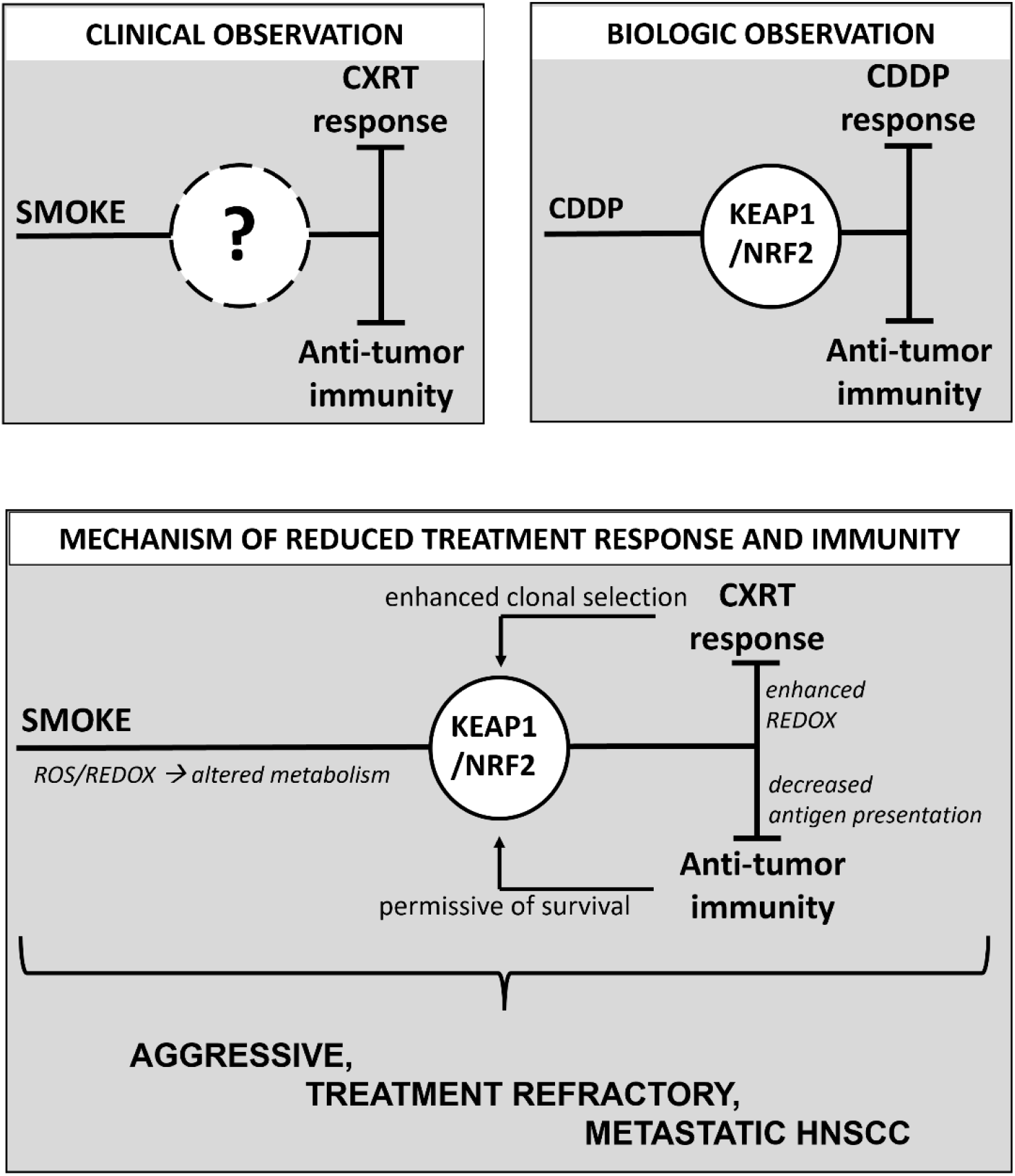
Mechanistic model of reduced treatment response and immunity. Clinical studies have definitively linked tobacco smoke exposure to reduced chemo-radiation (CXRT) response and altered immunity in HNSCC. Pre-clinical models of HNSCC have shown that hyper-activation of the Nrf2 pathway can drive resistance to cisplatin (CDDP) and alter the inflammatory profile of HNSCC cells and tumors. The current work proposes that the exposome, specifically smoke exposure activates a self-re-inforcing loop whereby activation of Nrf2 predisposes HNSCC tumors to reduced treatment response and suppressed immunity, generating an aggressive, treatment refractory and highly metastatic phenotype.

Over the last two decades, we and others have conclusively shown that smokers, particularly active smokers demonstrate inferior responses to chemoradiotherapy, regardless of HPV status, resulting in higher rates of disease recurrence and cancer specific death (3), (14). In parallel, we measured a distinct tumor immune microenvironment using overlapping approaches, strongly suggestive of suppressed anti-tumor immunity, presumably explaining a more aggressive clinical phenotype (**Fig.6**)(44). The current study sought to bring together these two observations into one comprehensive model, congruent with clinical reality. The model, outlined in **Fig.6** highlights an iterative process by which the smoke exposome predisposes a subset of HNSCC tumors to hyperactivation of the Nrf2 signaling pathway which is then exacerbated by initiation of chemoradiotherapy. Persistent and successful Nrf2 hyperactivation can then drive a dual phenotype of enhanced survival under conditions of ROS mediated stress through metabolic reprogramming and an altered secretory phenotype which can generate immune-suppressive effects.

As shown here, Keap1-Nrf2, although an exquisitely sensitive ROS sensor, and an adaptive mechanism for treatment resistance, can be maintained in a relatively narrow range as demonstrated by the difficulties associated with exogenous molecular manipulation of either *KEAP1* or *NFE2L2* (in contrast to routine manipulation of other oncogenes and tumor suppressors such as *TP53, PIK3CA, NOTCH1*). Together with evidence that the activation pattern of Nrf2 is rather short-lived but can be re-activated by repeated exposure, we can now understand a manner by which this pathway is persistently activated through chronic, daily exposure to cigarette smoke (and other environment toxins) leading to selection for cells and tumors predisposed toward chemoradiotherapy resistance. That persistent smoke exposure can directly lead to acquired cisplatin resistance is shown here directly as is the cross-resistance of cisplatin resistant cells to the oxidative stress generated by cigarette smoke. Notably, although e-cigarette vapor can deliver substantial nicotine levels to HNSCC cells, in contrast to conventional cigarette smoke, the oxidative stress component and the metabolic disruptions associated with the reductive state appear to be significantly diminished if not completely absent. Not only are cigarette smoke effects on activation of the Nrf2 pathway consistent across HNSCC cell lines of variable genomic background and HPV status, but the effects are clearly coordinated at transcriptional, translational and metabolic levels, with overall organization of the response aimed at enhancing the response to exogenous ROS.

Nrf2-hyperactivated human tumors have been shown by us and others to demonstrate an altered tumor immune microenvironment (TIME), with significant indications of suppressed, or inactive anti-tumor immunity.(39) Existing TCGA data clearly show a strong correlation between tobacco exposure, Nrf2 activation, and altered immunity. Here we present a rather comprehensive mechanistic analysis of this interaction and demonstrate that exposure of HNSCC to tobacco changes PDL1 levels. However, the combined impact on Nrf2 and NF-κB is more nuanced, and likely reflects the balance of cellular reducing potential. It is widely known that ROS triggers both Nrf2 and NF-kB, which have antagonistic functions.(45) Elevated ROS activates NF-kB activity leading to increased inflammatory mediators, whereas downstream targets of Nrf2 reduce ROS to potentially inhibit NF-κB and shut down inflammation. Since both pathways are simultaneously triggered by cigarette smoke, the final phenotype depends on their balance which is intimately linked to exposure levels, timing, and redox homeostasis―all of which can be difficult to experimentally model. In light of our previous work connecting Nrf2 activation to an immunosuppressive tumor immune microenvironment for smoking related cancers, it is likely that this pathway dominates these tumors over the long run. Regardless, there is a clear connection between cellular insults and the inflammatory secretome of tumors. This was apparent by changes in the inflammatory mediators PGE2 and IL-6 secreted by smoke exposed cells, through experiments showing conditioned media from smoke exposed tumor cells can rewire PBMC programing and differentiation, and by changes in the tumor immune microenvironment we found *in vivo* with smoke exposure.

Our data support the translational conclusion that Nrf2 activation represents a functional nexus between metabolic reprogramming designed to support oxidative stress, development of treatment resistance and altered immunity. Several questions remain unanswered. First, what is the role of nicotine, exclusive of oxidative stress? Nicotine has been shown to stimulate cancer progression mediated by the activation of nicotinic acetylcholine receptors (nAChRs) (46). This signaling has been linked to promoting lung carcinogenesis and facilitating immune evasion by smoke-induced macrophages in NSCLC cells (47). Similar research also suggests that activating nAChR7 contributes to immune evasion triggered by cigarette smoke, by increasing PD-L1 expression levels in lung epithelial cells (48). Although lacking an oxidative stress component, e-cigarette vapor exposure will require more extensive *in vivo* modelling to determine whether the immunomodulatory effects we capture with conventional cigarette smoke are at least partially replicated in the absence of the oxidative volatile agents.

Second, although we have identified what we believe are the critical components of the Nrf2 pathway related to oxidative stress response and altered metabolism both here and in our previous publications (15), (16), (18), focused screens using either shRNA or CRISPR will be required to determine how many down-stream effectors are both necessary *and* sufficient to support the reductive state adaptation described here, the acquisition of chemoradiotherapy resistance, and altered immunity. Finally, newly developed GEMM models, which are only partially leveraged here for their stable Keap1 suppressed status, will require further examination to better understand the effects of chronic cigarette smoke exposure and development of chemoradiotherapy sensitivity and/or resistance.

## References

1. Johnson DE, Burtness B, Leemans CR, Lui VWY, Bauman JE, Grandis JR. Head and neck squamous cell carcinoma. Nature Reviews Disease Primers 2020;6(1):92 doi 10.1038/s41572-020-00224-3.

2. Spiotto MT, Taniguchi CM, Klopp AH, Colbert LE, Lin SH, Wang L, et al. Biology of the Radio- and Chemo-Responsiveness in HPV Malignancies. Semin Radiat Oncol 2021;31(4):274–85 doi 10.1016/j.semradonc.2021.02.009.

3. Elhalawani H, Mohamed ASR, Elgohari B, Lin TA, Sikora AG, Lai SY, et al. Tobacco exposure as a major modifier of oncologic outcomes in human papillomavirus (HPV) associated oropharyngeal squamous cell carcinoma. BMC Cancer 2020;20(1):912 doi 10.1186/s12885-020-07427-7.

4. Sandulache VC, Wilde DC, Sturgis EM, Chiao EY, Sikora AG. A Hidden Epidemic of “Intermediate Risk” Oropharynx Cancer. Laryngoscope Investig Otolaryngol 2019;4(6):617–23 doi 10.1002/lio2.316.

5. Jethwa AR, Khariwala SS. Tobacco-related carcinogenesis in head and neck cancer. Cancer Metastasis Rev 2017;36(3):411–23 doi 10.1007/s10555-017-9689-6.

6. van Imhoff LC, Kranenburg GG, Macco S, Nijman NL, van Overbeeke EJ, Wegner I, et al. Prognostic value of continued smoking on survival and recurrence rates in patients with head and neck cancer: A systematic review. Head Neck 2016;38 Suppl 1:E2214-20 doi 10.1002/hed.24082.

7. Wahle BM, Zolkind P, Ramirez RJ, Skidmore ZL, Anderson SR, Mazul A, et al. Integrative genomic analysis reveals low T-cell infiltration as the primary feature of tobacco use in HPV-positive oropharyngeal cancer. iScience 2022;25(5):104216 doi 10.1016/j.isci.2022.104216.

8. Liang F, Wang GZ, Wang Y, Yang YN, Wen ZS, Chen DN, et al. Tobacco carcinogen induces tryptophan metabolism and immune suppression via induction of indoleamine 2,3-dioxygenase 1. Signal Transduct Target Ther 2022;7(1):311 doi 10.1038/s41392-022-01127-3.

9. Guo J, Kim D, Gao J, Kurtyka C, Chen H, Yu C, et al. IKBKE is induced by STAT3 and tobacco carcinogen and determines chemosensitivity in non-small cell lung cancer. Oncogene 2013;32(2):151–9 doi 10.1038/onc.2012.39.

10. Caliri AW, Tommasi S, Besaratinia A. Relationships among smoking, oxidative stress, inflammation, macromolecular damage, and cancer. Mutat Res Rev Mutat Res 2021;787:108365 doi 10.1016/j.mrrev.2021.108365.

11. Wang R, Liang L, Matsumoto M, Iwata K, Umemura A, He F. Reactive Oxygen Species and NRF2 Signaling, Friends or Foes in Cancer? Biomolecules 2023;13(2) doi 10.3390/biom13020353.

12. Aggarwal V, Tuli HS, Varol A, Thakral F, Yerer MB, Sak K, et al. Role of Reactive Oxygen Species in Cancer Progression: Molecular Mechanisms and Recent Advancements. Biomolecules 2019;9(11) doi 10.3390/biom9110735.

13. Castle PE. How does tobacco smoke contribute to cervical carcinogenesis? J Virol 2008;82(12):6084-5; author reply 5-6 doi 10.1128/JVI.00103-08.

14. Wilde DC, Castro PD, Bera K, Lai S, Madabhushi A, Corredor G, et al. Oropharyngeal cancer outcomes correlate with p16 status, multinucleation and immune infiltration. Mod Pathol 2022;35(8):1045–54 doi 10.1038/s41379-022-01024-8.

15. Osman AA, Arslan E, Bartels M, Michikawa C, Lindemann A, Tomczak K, et al. Dysregulation and Epigenetic Reprogramming of NRF2 Signaling Axis Promote Acquisition of Cisplatin Resistance and Metastasis in Head and Neck Squamous Cell Carcinoma. Clin Cancer Res 2023;29(7):1344–59 doi 10.1158/1078-0432.CCR-22-2747.

16. Yu W, Chen Y, Putluri N, Osman A, Coarfa C, Putluri V, et al. Evolution of cisplatin resistance through coordinated metabolic reprogramming of the cellular reductive state. Br J Cancer 2023;128(11):2013–24 doi 10.1038/s41416-023-02253-7.

17. Ahmed KM, Veeramachaneni R, Deng D, Putluri N, Putluri V, Cardenas MF, et al. Glutathione peroxidase 2 is a metabolic driver of the tumor immune microenvironment and immune checkpoint inhibitor response. J Immunother Cancer 2022;10(8) doi 10.1136/jitc-2022-004752.

18. Yu W, Chen Y, Putluri N, Coarfa C, Robertson MJ, Putluri V, et al. Acquisition of Cisplatin Resistance Shifts Head and Neck Squamous Cell Carcinoma Metabolism toward Neutralization of Oxidative Stress. Cancers (Basel) 2020;12(6) doi 10.3390/cancers12061670.

19. Frederick M, Skinner HD, Kazi SA, Sikora AG, Sandulache VC. High expression of oxidative phosphorylation genes predicts improved survival in squamous cell carcinomas of the head and neck and lung. Sci Rep 2020;10(1):6380 doi 10.1038/s41598-020-63448-z.

20. Mertins P, Yang F, Liu T, Mani DR, Petyuk VA, Gillette MA, et al. Ischemia in tumors induces early and sustained phosphorylation changes in stress kinase pathways but does not affect global protein levels. Mol Cell Proteomics 2014;13(7):1690–704 doi 10.1074/mcp.M113.036392.

21. Sutton C, Nozawa K, Kent K, Saltzman A, Leng M, Nagarajan S, et al. Molecular dissection and testing of PRSS37 function through LC-MS/MS and the generation of a PRSS37 humanized mouse model. Sci Rep 2023;13(1):11374 doi 10.1038/s41598-023-37700-1.

22. da Veiga Leprevost F, Haynes SE, Avtonomov DM, Chang HY, Shanmugam AK, Mellacheruvu D, et al. Philosopher: a versatile toolkit for shotgun proteomics data analysis. Nat Methods 2020;17(9):869–70 doi 10.1038/s41592-020-0912-y.

23. Monroe ME, Shaw JL, Daly DS, Adkins JN, Smith RD. MASIC: a software program for fast quantitation and flexible visualization of chromatographic profiles from detected LC-MS(/MS) features. Comput Biol Chem 2008;32(3):215–7 doi 10.1016/j.compbiolchem.2008.02.006.

24. Kong AT, Leprevost FV, Avtonomov DM, Mellacheruvu D, Nesvizhskii AI. MSFragger: ultrafast and comprehensive peptide identification in mass spectrometry-based proteomics. Nat Methods 2017;14(5):513–20 doi 10.1038/nmeth.4256.

25. Fondrie WE, Noble WS. mokapot: Fast and Flexible Semisupervised Learning for Peptide Detection. J Proteome Res 2021;20(4):1966–71 doi 10.1021/acs.jproteome.0c01010.

26. Saltzman AB, Leng M, Bhatt B, Singh P, Chan DW, Dobrolecki L, et al. gpGrouper: A Peptide Grouping Algorithm for Gene-Centric Inference and Quantitation of Bottom-Up Proteomics Data. Mol Cell Proteomics 2018;17(11):2270–83 doi 10.1074/mcp.TIR118.000850.

27. Gu Z, Eils R, Schlesner M. Complex heatmaps reveal patterns and correlations in multidimensional genomic data. Bioinformatics 2016;32(18):2847–9 doi 10.1093/bioinformatics/btw313.

28. Kami Reddy KR, Piyarathna DWB, Park JH, Putluri V, Amara CS, Kamal AHM, et al. Mitochondrial reprogramming by activating OXPHOS via glutamine metabolism in African American patients with bladder cancer. JCI Insight 2024;9(17) doi 10.1172/jci.insight.172336.

29. Vantaku V, Putluri V, Bader DA, Maity S, Ma J, Arnold JM, et al. Epigenetic loss of AOX1 expression via EZH2 leads to metabolic deregulations and promotes bladder cancer progression. Oncogene 2020;39(40):6265–85 doi 10.1038/s41388-019-0902-7.

30. Wishart DS, Tzur D, Knox C, Eisner R, Guo AC, Young N, et al. HMDB: the Human Metabolome Database. Nucleic Acids Res 2007;35(Database issue):D521-6 doi 10.1093/nar/gkl923.

31. Piyarathna DWB, Rajendiran TM, Putluri V, Vantaku V, Soni T, von Rundstedt FC, et al. Distinct Lipidomic Landscapes Associated with Clinical Stages of Urothelial Cancer of the Bladder. Eur Urol Focus 2018;4(6):907–15 doi 10.1016/j.euf.2017.04.005.

32. Vantaku V, Dong J, Ambati CR, Perera D, Donepudi SR, Amara CS, et al. Multi-omics Integration Analysis Robustly Predicts High-Grade Patient Survival and Identifies CPT1B Effect on Fatty Acid Metabolism in Bladder Cancer. Clin Cancer Res 2019;25(12):3689–701 doi 10.1158/1078-0432.CCR-18-1515.

33. Liberzon A, Birger C, Thorvaldsdottir H, Ghandi M, Mesirov JP, Tamayo P. The Molecular Signatures Database (MSigDB) hallmark gene set collection. Cell Syst 2015;1(6):417–25 doi 10.1016/j.cels.2015.12.004.

34. Skinner HD, Sandulache VC, Ow TJ, Meyn RE, Yordy JS, Beadle BM, et al. TP53 disruptive mutations lead to head and neck cancer treatment failure through inhibition of radiation-induced senescence. Clin Cancer Res 2012;18(1):290–300 doi 10.1158/1078-0432.CCR-11-2260.

35. Sandulache VC, Ow TJ, Pickering CR, Frederick MJ, Zhou G, Fokt I, et al. Glucose, not glutamine, is the dominant energy source required for proliferation and survival of head and neck squamous carcinoma cells. Cancer 2011;117(13):2926–38 doi 10.1002/cncr.25868.

36. Hudson WH, Olson JJ, Sudmeier LJ. Immune microenvironment remodeling after radiation of a progressing brain metastasis. Cell Rep Med 2023;4(6):101054 doi 10.1016/j.xcrm.2023.101054.

37. Singh A, Rangasamy T, Thimmulappa RK, Lee H, Osburn WO, Brigelius-Flohe R, et al. Glutathione peroxidase 2, the major cigarette smoke-inducible isoform of GPX in lungs, is regulated by Nrf2. Am J Respir Cell Mol Biol 2006;35(6):639–50 doi 10.1165/rcmb.2005-0325OC.

38. Li X, Ye Y, Peng K, Zeng Z, Chen L, Zeng Y. Histones: The critical players in innate immunity. Front Immunol 2022;13:1030610 doi 10.3389/fimmu.2022.1030610.

39. Kazi MA, Veeramachaneni R, Deng D, Putluri N, Cardinas M, Sikora A, et al. Glutathione peroxidase 2 is a metabolic driver of the tumor immune microenvironment and immune checkpoint inhibitor response. Journal for ImmunoTherapy of Cancer 2022.

40. Veeramachaneni R, Yu W, Newton JM, Kemnade JO, Skinner HD, Sikora AG, et al. Metformin generates profound alterations in systemic and tumor immunity with associated antitumor effects. J Immunother Cancer 2021;9(7) doi 10.1136/jitc-2021-002773.

41. Nakamura H, Takada K. Reactive oxygen species in cancer: Current findings and future directions. Cancer Sci 2021;112(10):3945–52 doi 10.1111/cas.15068.

42. Taguchi K, Yamamoto M. The KEAP1-NRF2 System as a Molecular Target of Cancer Treatment. Cancers (Basel) 2020;13(1) doi 10.3390/cancers13010046.

43. Dhawan A, Pifer PM, Sandulache VC, Skinner HD. Metabolic targeting, immunotherapy and radiation in locally advanced non-small cell lung cancer: Where do we go from here? Front Oncol 2022;12:1016217 doi 10.3389/fonc.2022.1016217.

44. Kemnade JO, Elhalawani H, Castro P, Yu J, Lai S, Ittmann M, et al. CD8 infiltration is associated with disease control and tobacco exposure in intermediate-risk oropharyngeal cancer. Sci Rep 2020;10(1):243 doi 10.1038/s41598-019-57111-5.

45. Casper E. The crosstalk between Nrf2 and NF-kappaB pathways in coronary artery disease: Can it be regulated by SIRT6? Life Sci 2023;330:122007 doi 10.1016/j.lfs.2023.122007.

46. Schuller HM. Is cancer triggered by altered signalling of nicotinic acetylcholine receptors? Nat Rev Cancer 2009;9(3):195–205 doi 10.1038/nrc2590.

47. Kang G, Jiao Y, Pan P, Fan H, Li Q, Li X, et al. alpha5-nAChR/STAT3/CD47 axis contributed to nicotine-related lung adenocarcinoma progression and immune escape. Carcinogenesis 2023;44(10-11):773-84 doi 10.1093/carcin/bgad061.

48. Kwok HH, Gao B, Chan KH, Ip MS, Minna JD, Lam DC. Nicotinic Acetylcholine Receptor Subunit alpha7 Mediates Cigarette Smoke-Induced PD-L1 Expression in Human Bronchial Epithelial Cells. Cancers (Basel) 2021;13(21) doi 10.3390/cancers13215345.

