## Supplemental Figures for "Tobacco smoke exposure is a driver of altered oxidative stress response and immunity in head and neck cancer"

### SUPPLEMENTAL FIGURE 1

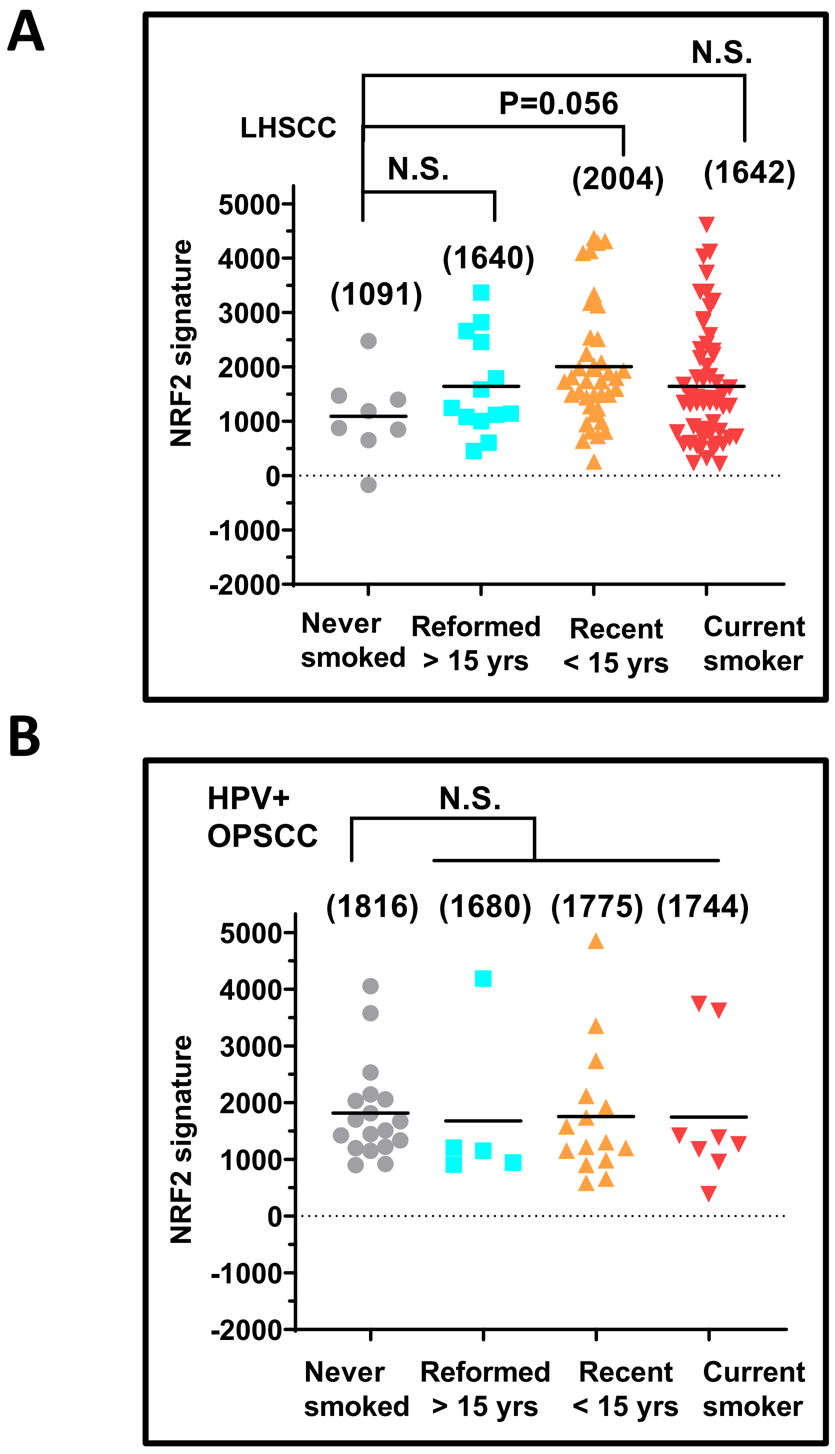


**Supplemental Figure 1. NRF2 activation and smoking history in other HNSCC subsites**. (A) A trend towards increased NRF2 activation was found in LHSCC TCGA tumors from patients with a smoking history. (B) NRF2 activation is increased in HPV-associated OPSCC (HPV-associated), independent of smoking history.

#### SUPPLEMENTAL FIGURE 2

A

B

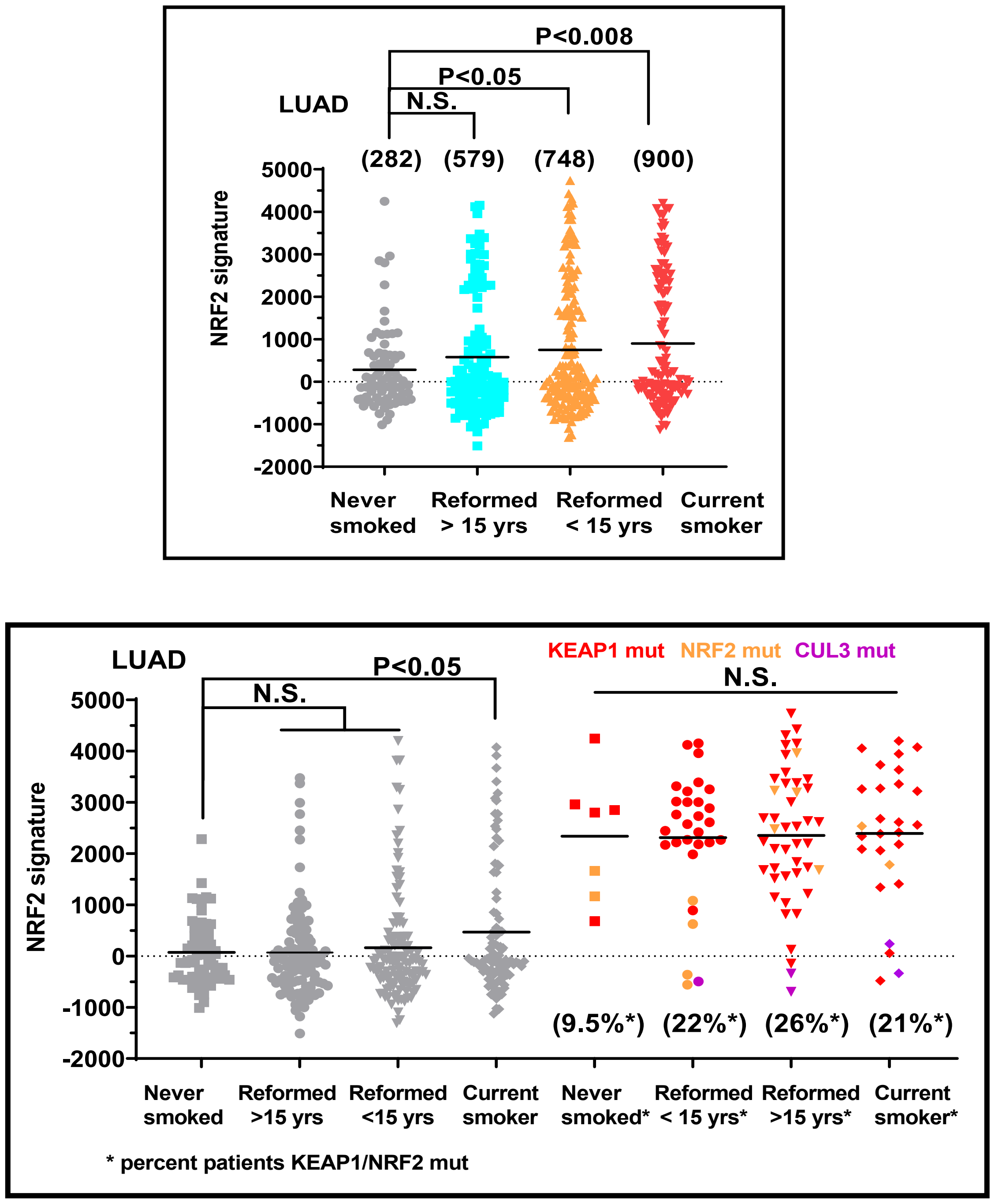


| | Never smoker | Smoker |
|---|---|---|
| **Wild type** | 67 (58) | 323 (331) |
| **KEAP1/NRF2 mutation** | 7 (16) | 97 (88) |
| P= 0.012 | | |

**Supplemental Figure 2. NRF2 activation is elevated in LUAD tumors from patients with a recent history of smoking.** A) NRF2 activation scores are significantly elevated in tumors from patients who currently smoke or recently quit compared to never-smokers. B) Among LUAD tumors wild type for KEAP1/NRF2, NRF2 activity remains elevated in current smokers, while the prevalence of KEAP1/NRF2 mutations is lower in never smokers compared to patients who have some history of smoking where the frequency is more than doubled. The association between mutation frequency and smoking history was significant (P=0.012) by a Fisher's exact test.

#### SUPPLEMENTAL FIGURE 3

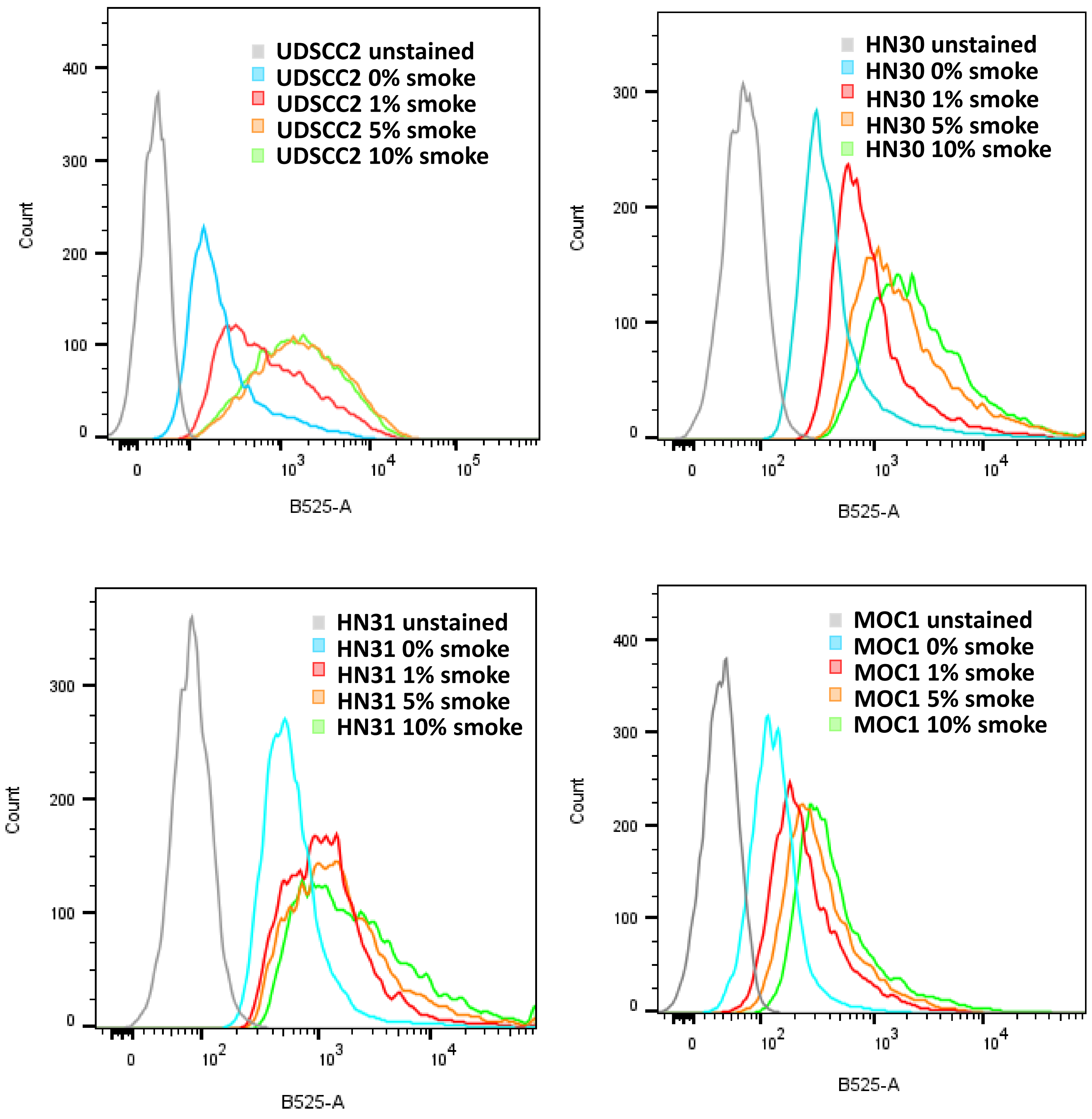


**Supplemental Figure 3. Tobacco exposure effects are ROS-mediated.** Cellular ROS levels were detected by measuring the fluorescence of DCFH-DA (B525-A) in UDSCC2, HN30, HN31, and MOC1 cells after treating cells with various concentrations of smoke-infused media for 6 h.

**SUPPLEMENTAL FIGURE 4**

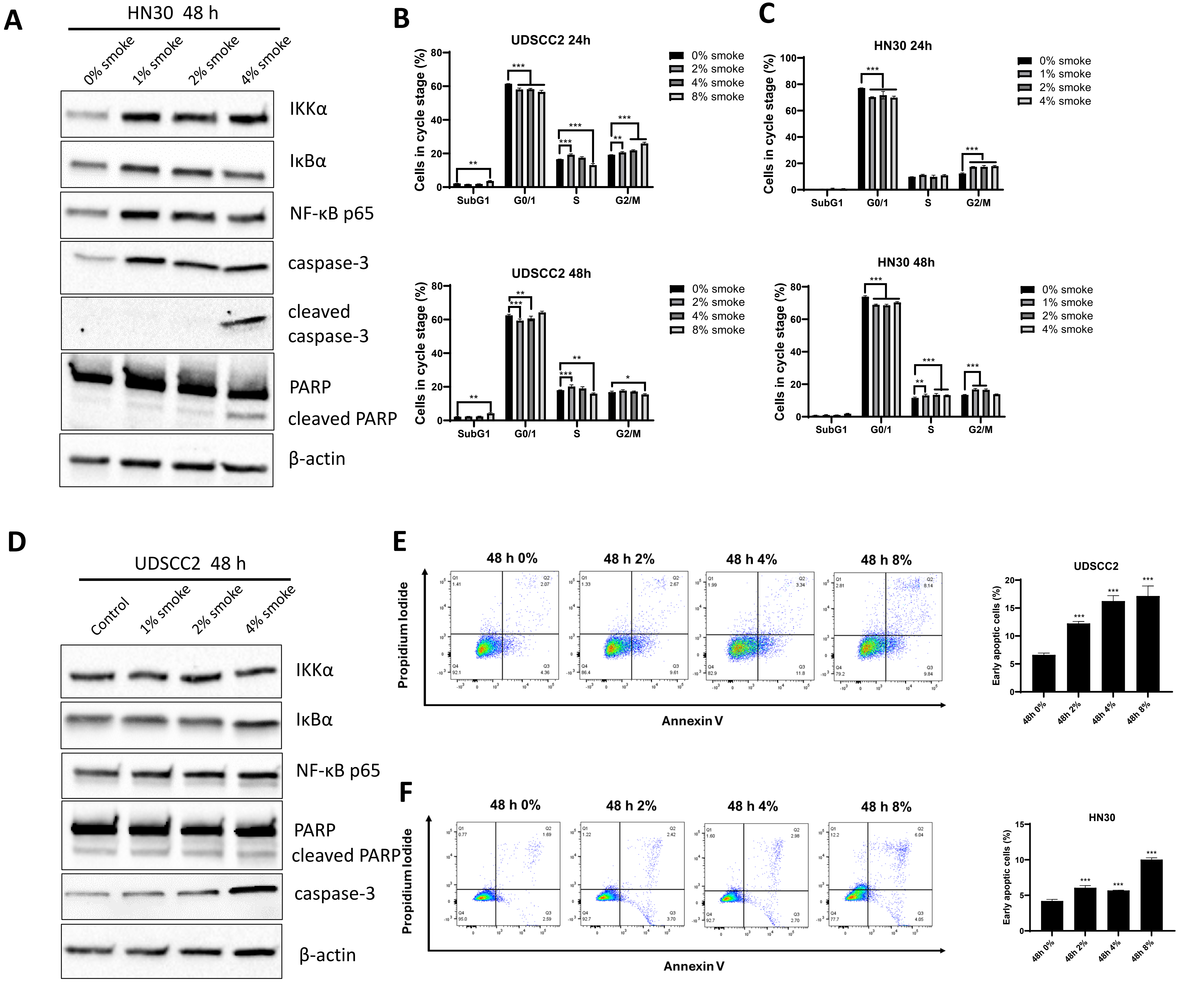


**Supplemental Figure 4. Tobacco exposure induces cell death in HNSCC.** Western blot analysis of HN30 (A) and UDSCC2 (D) cells at 48 hours post-smoke exposure was conducted to assess NFkB activation and the cleavage of PARP and caspase-3. Propidium iodide (PI) staining was employed to evaluate the impact of smoke on cell cycle progression in HN30 (B) and UDSCC2 (C) cells. PI combined with Annexin V staining was performed to examine the effects of smoke on early apoptosis of UDSCC2 (E) and HN30 (F) cells. *p<0.05, **P<0.05, and ***p<0.001

#### SUPPLEMENTAL FIGURE 5

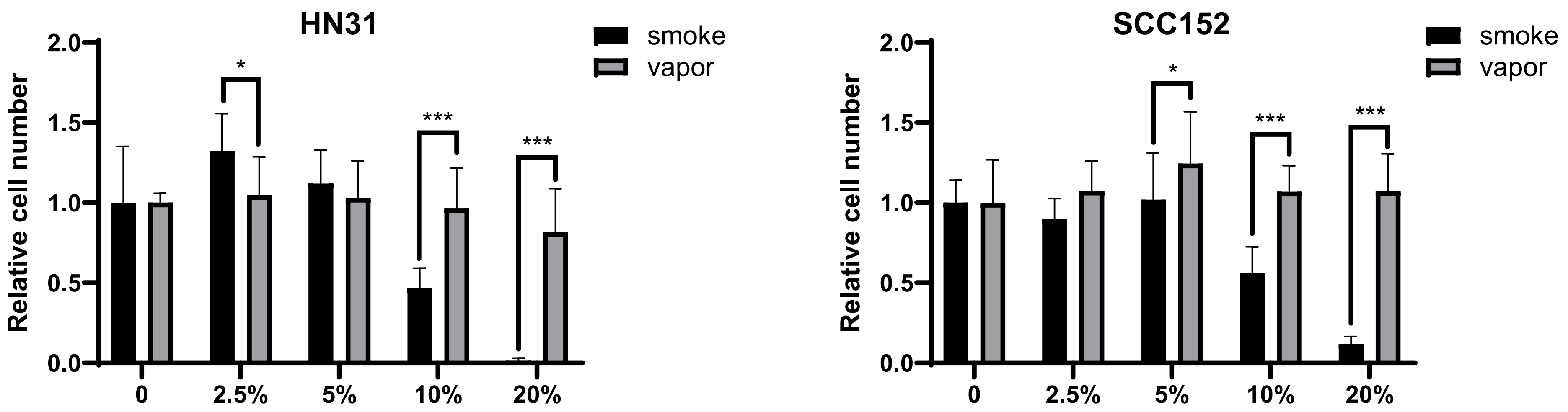


**Supplemental Figure 5. Comparison of cytotoxicity between cigarette smoke and E-cigarette smoke in HPV-Independent (HN31) and HPV-Associated (SCC152) HNSCC cells.** HN31 and SCC152 cells were exposed to media infused with 0%, 2.5%, 5%, 10%, and 20% cigarette or e-cigarette smoke concentrations. The relative cell numbers were assessed using the Hoechst assay after a 72-hour exposure period. Statistical significance was denoted as *p<0.05, **p<0.01, ***p<0.001 for comparisons between values.

#### SUPPLEMENTAL FIGURE 6

A

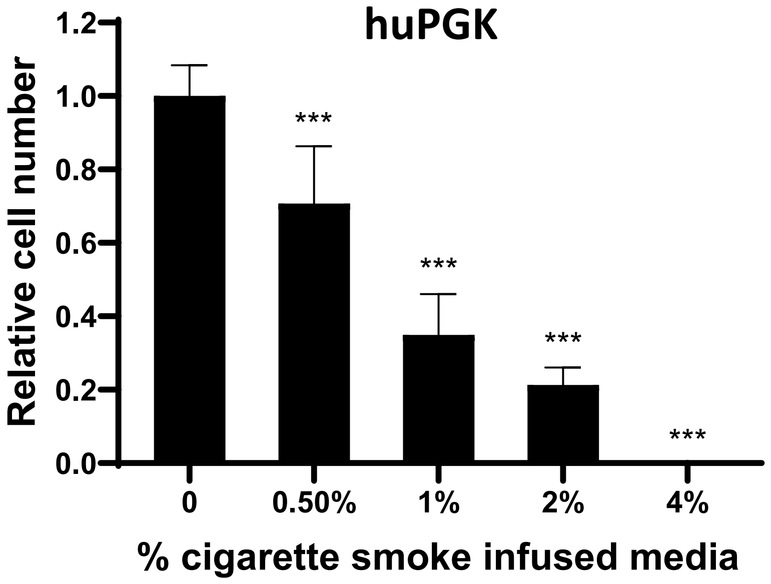


B

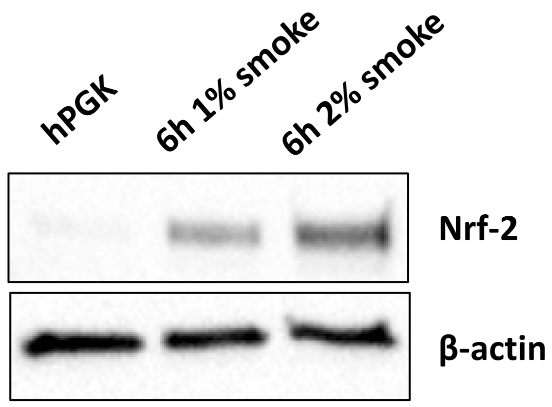


**Supplemental Figure 6. Effect of smoke exposure in human primary gingival keratinocytes cells**. (A) Human primary gingival keratinocytes were exposed to 0.5%, 1%, 2%, and 4% concentrations of cigarette smoke-infused media. The relative cell numbers were assessed using the Hoechst assay after a 72-hour exposure period. (B) Human primary gingival keratinocytes were exposed to 1%, and 2% smoke media for 6 h before harvesting for Nrf2 protein expression. *p<0.05, **P<0.05, and ***p<0.001

### SUPPLEMENTAL FIGURE 7

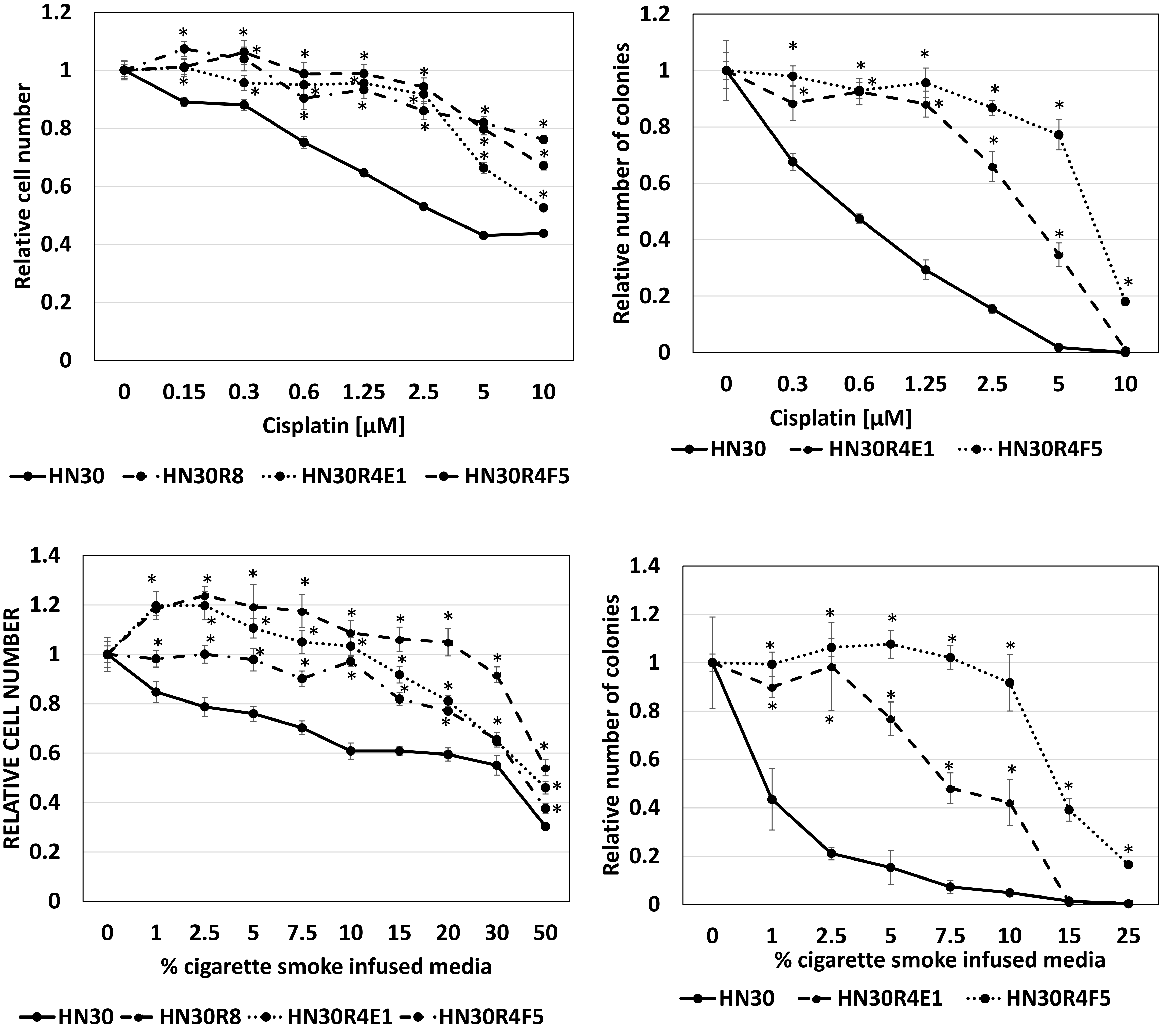


**Supplemental Figure 7. Cisplatin – smoke cross resistance.** Using a conditioned resistance model, we generated individual clones of HN30 which demonstrate acquired cisplatin resistance. Using a 72-hour Hoechst assay we compared the cell viability of cisplatin (upper panel) or cigarette smoke exposure (lower panel) in the parental HN30 cell line and its cisplatin-resistant clones HN30R8, HN30R4E1, and HN30R4F5. Data are shown as means, normalized to a control condition; error bars indicate standard deviation. Statistically significant deviations were observed from parental HN30 vs. cisplatin-resistant clones HN30R8, HN30R4E1, and HN30R4F5 for each treatment (*$p < 0.05$).

#### SUPPLEMENTAL FIGURE 8

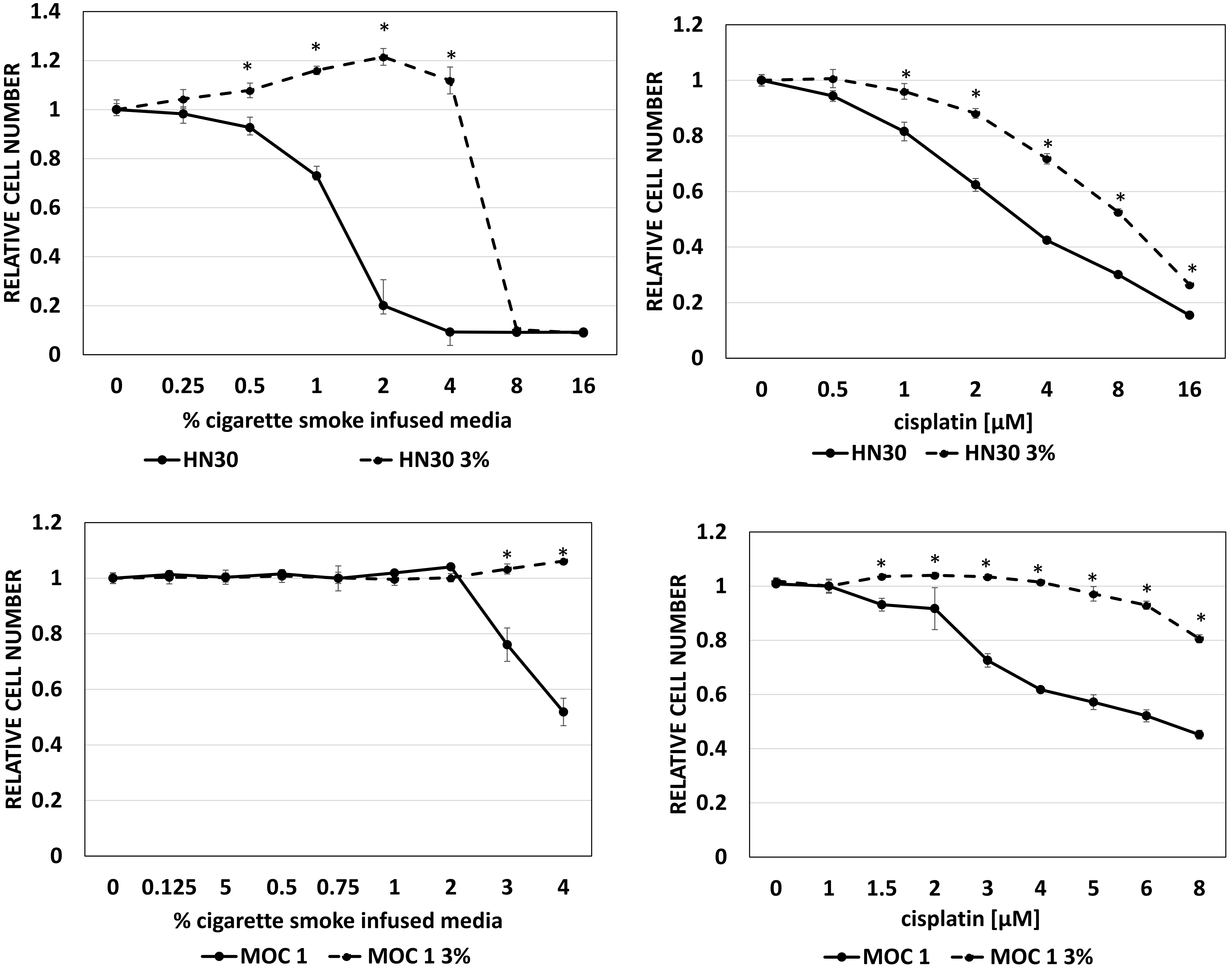


**Supplemental Figure 8. Smoke – cisplatin cross resistance.** A 72-hour Hoechst assay was used to compare the cell viability of cigarette smoke infused media and cisplatin treatment in the parental cell lines, and their corresponding 3% chronically exposure models (HN30 in the upper panel and MOC1 cells in the lower panel). Data are shown as means, normalized to a control condition; error bars indicate standard deviation. Statistically significant deviations were observed from parental HN30 vs. HN30 3% and MOC1 vs. MOC1 3% for each treatment (*$p < 0.05$).

#### SUPPLEMENTAL FIGURE 9

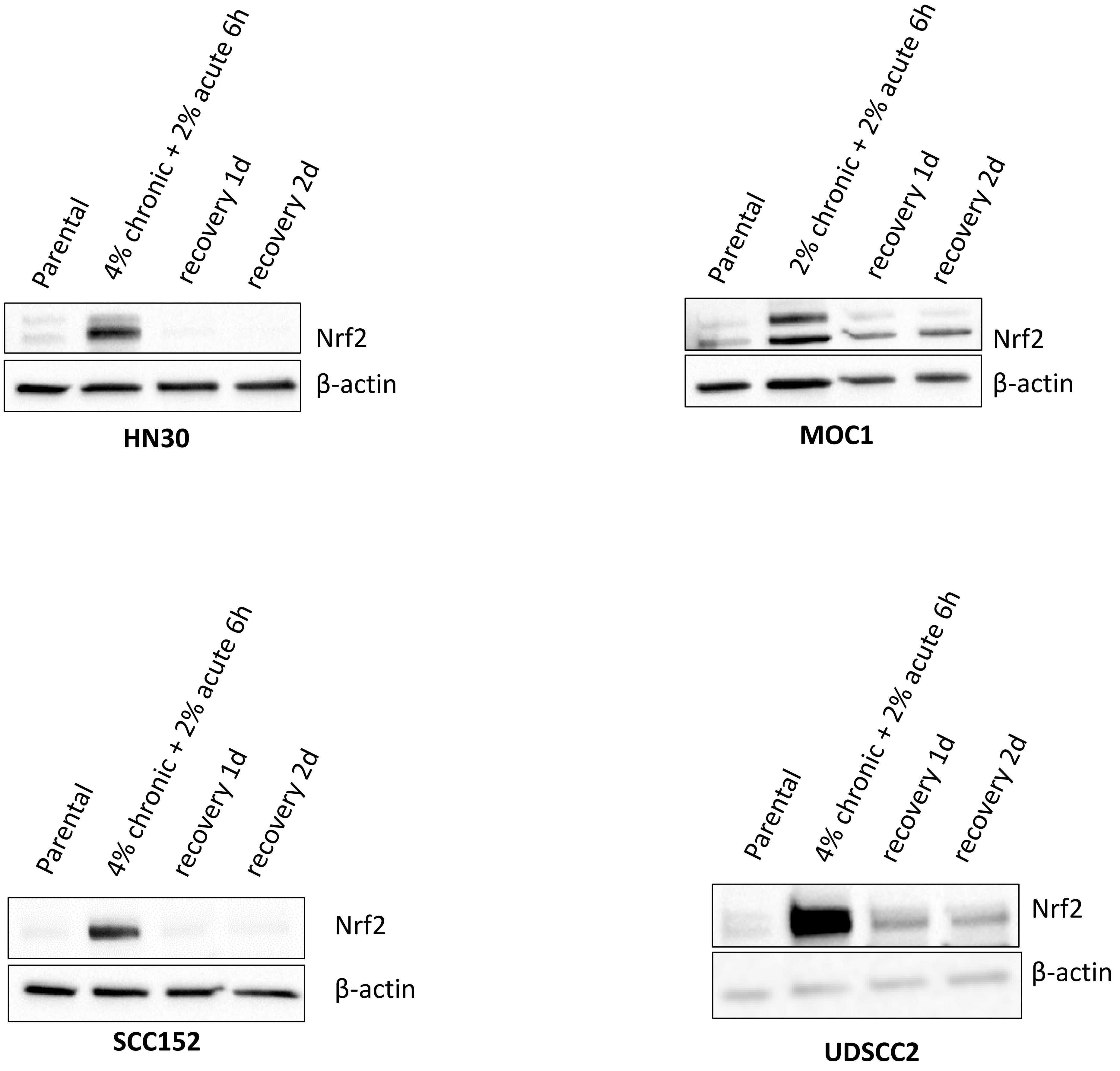


**Supplemental Figure 9. Transient Nrf2 stabilization.** A chronically smoke exposed HPV-independent HNSCC cell line (HN30), an HPV-associated HNSCC cell line (SCC152), and a murine OCSCC model (MOC1) were treated with an additional bolus of 6 h of the cigarette infused media. After removing the infused smoke media, the cells were given a 1 and 2 days to recover before harvesting. An HPV-associated HNSCC cell line (UDSCC2) was chronically exposed to cigarette-infused media at the indicated concentrations (0.75%, 1%, and 2%) for several months followed by an additional bolus for 6 and 24 h. Cells were harvested for Nrf-2 expression by western blot analysis and β-actin as the protein loading control.

#### SUPPLEMENTAL FIGURE 10

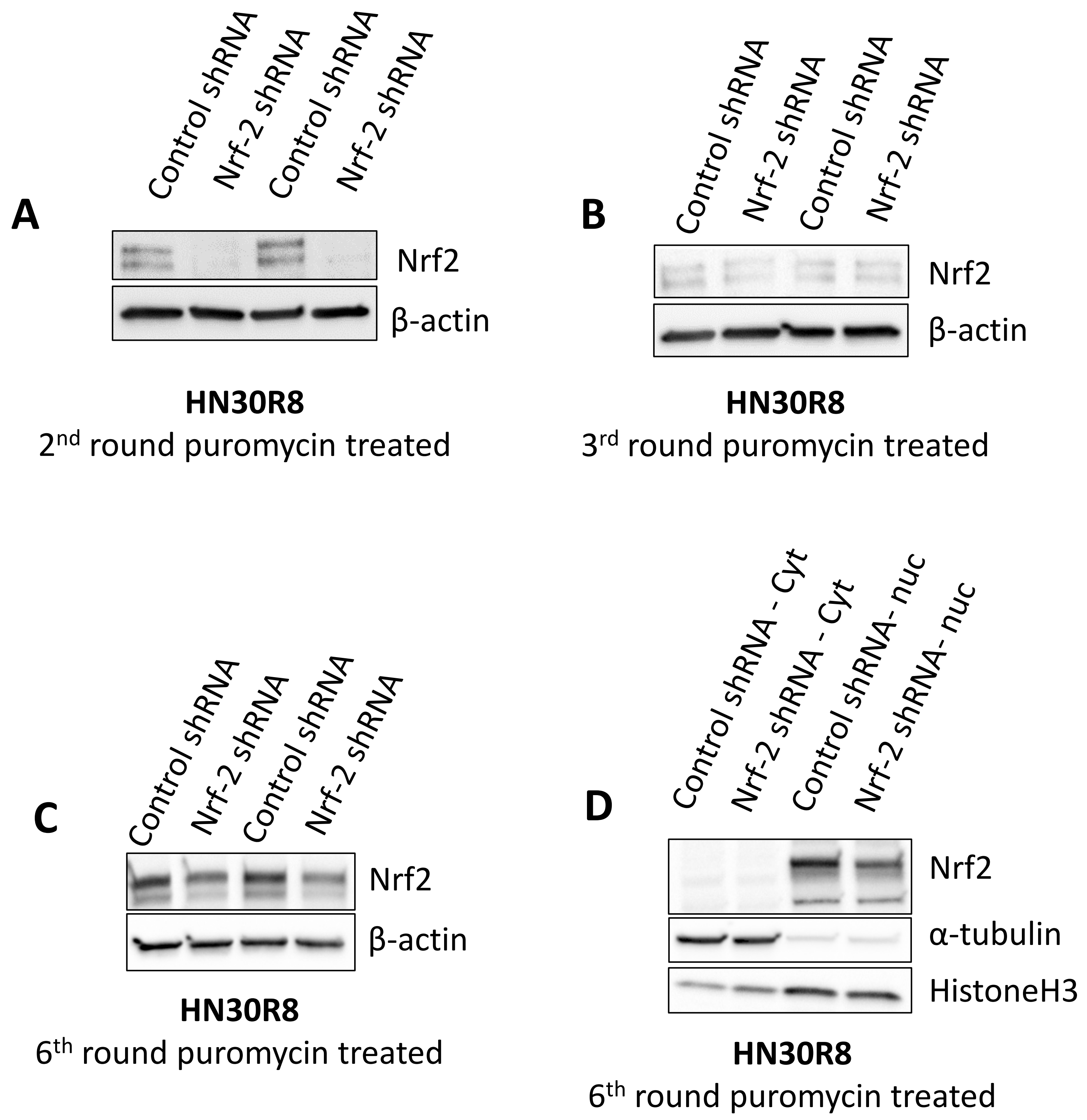


**Supplemental Figure 10. Unstable Nrf2 manipulation.** HN30R8 cells were infected with lentivirus constructs containing either control vector or shRNA targeting Nrf-2 from Santa Cruz. Cells were harvested after $2^{nd}$ (A), $3^{rd}$ (B), and $6^{th}$ (C) round puromycin selection, and the expression of Nrf-2 protein were detected by western blot. (D) The levels of Nrf-2 in cell nuclear and cytoplasm extracts of HN30R8 were analyzed. α-Tubulin and Histone H3 served as loading and purity controls for the cytoplasm and nuclear fractions, respectively.

### SUPPLEMENTAL FIGURE 11

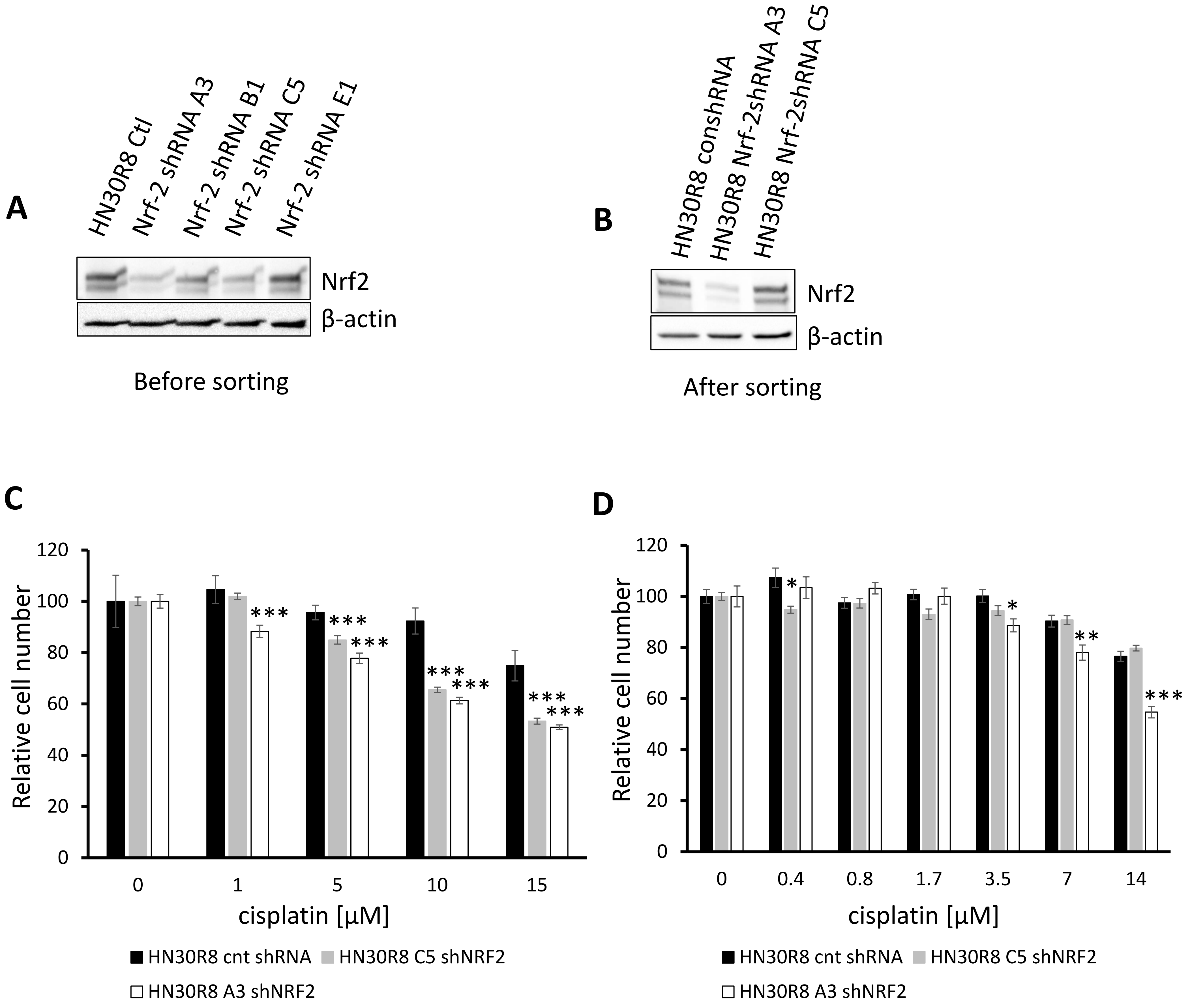


**Supplemental Figure 11. Unstable Nrf2 manipulation and phenotype.** HEK293T cells were transfected with either four different shRNA plasmids targeting Nrf-2 (A3, B1, C5, E1) or control vector from GIPZ. The corresponding viruses were collected and infected with HN30R8 cells. After 4ug/mL puromycin treatment for 3 days, cells were harvested for whole cell lysate, followed by Western blot analysis for Nrf-2 knockdown (A). (B) After several passages, cells were sorted for 40%-90% EGFP-positive cells to obtain highly enriched fractions of homogenously and strongly transfected cells. Nrf-2 Knockdown efficiency was less efficient compared to the unsorted cells. (C) Hoechst assay was performed to compare the sensitivity between the control shRNA vs. Nrf-2 shRNA before and after sorting (D) when cells were treated with various concentrations of cisplatin for 3 days. *p<0.05, **P<0.05, and ***p<0.001

#### SUPPLEMENTAL FIGURE 12

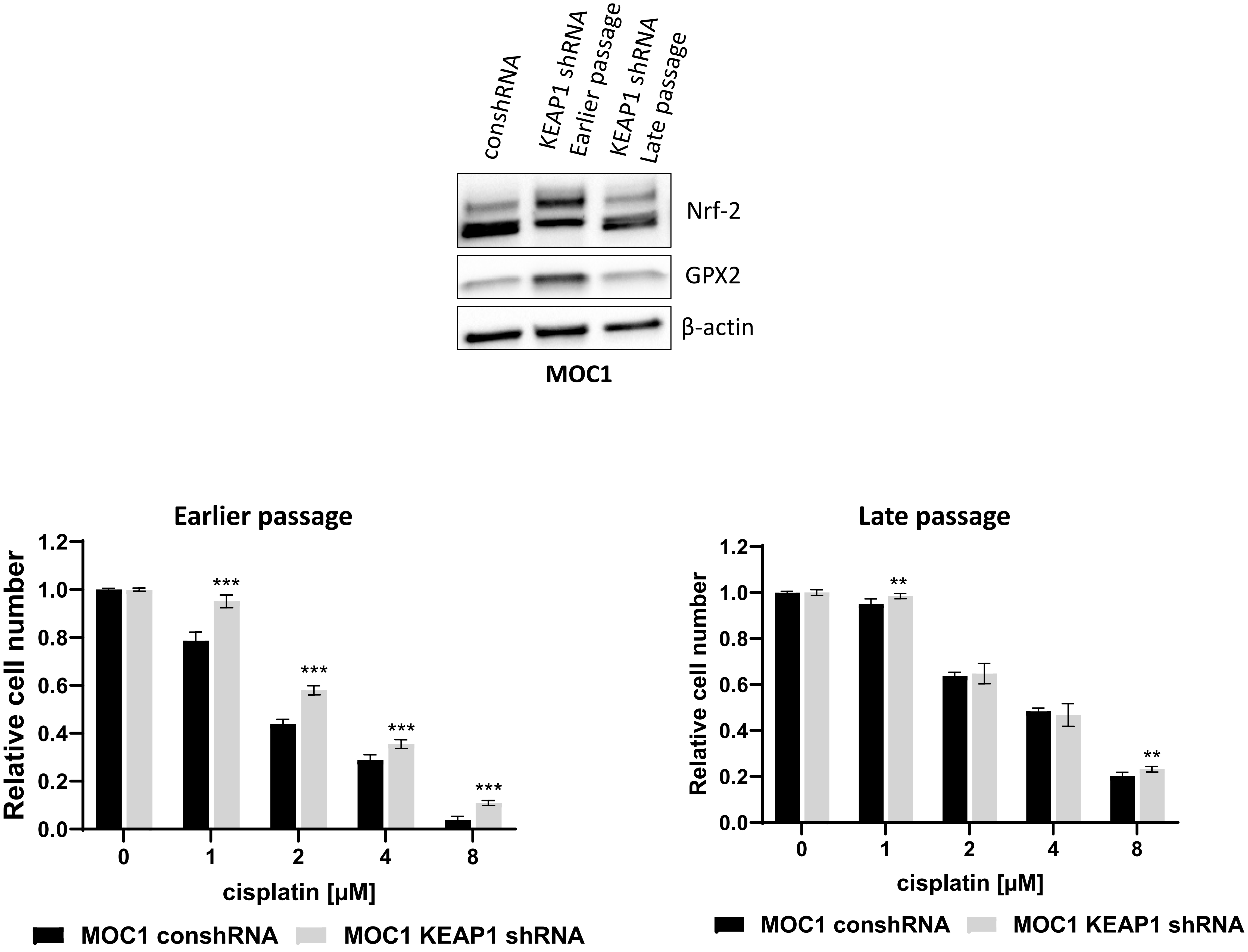


**Supplemental Figure 12. Unstable Keap1 manipulation and phenotype.** HEK293T cells were transfected with shRNA plasmids targeting Keap1 or control vector from GIPZ. The corresponding viruses were collected and infected with murine MOC1 cells. After 4ug/mL puromycin treatment for 3 days, cells were harvested for whole cell lysate, followed by Western blot analysis for Keap1. Cells were harvested immediately after the stable cell lines were established (earlier passage) and cultured after one month (late passage) for Nrf-2 and GPX2 protein expression. Hoechst assay was performed to compare the sensitivity between the control shRNA vs. Keap1 shRNA for the earlier and late passages that were treated with various concentrations of cisplatin for 3 days. *p<0.05, **P<0.05, and ***p<0.001

#### SUPPLEMENTAL FIGURE 13

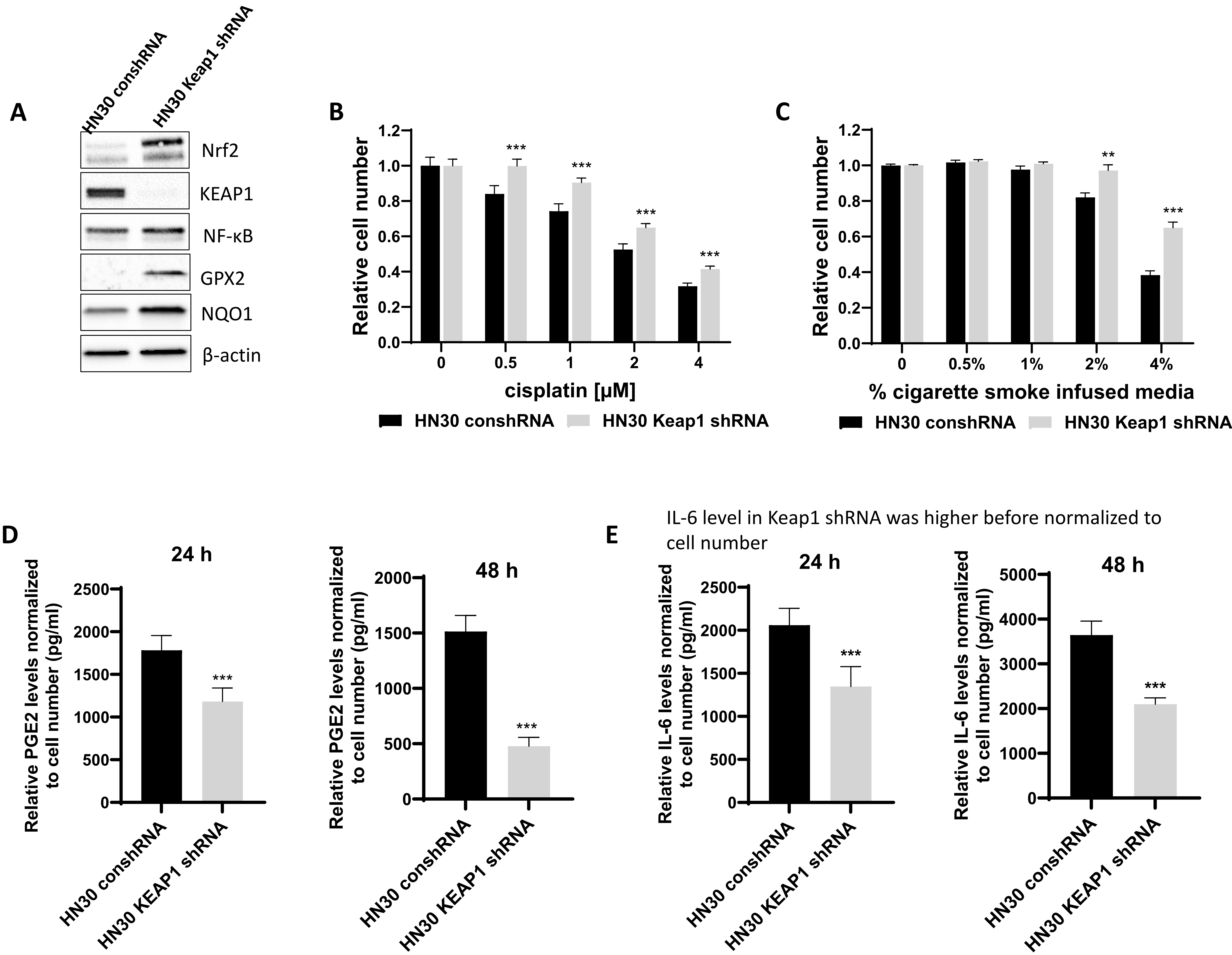


**Supplemental Figure 13. KEAP1 effects on PGE2 and IL-6 production.** HEK293T cells were transfected with either human keap1 shRNA plasmid or conshRNA from GIPZ. The corresponding viruses were collected and infected with HN30 (A) cells. After 4 μg/mL puromycin treatment for 3 days, cells were harvested for whole cell lysate, followed by Western blot analysis for the Nrf-2 targeted proteins. Hoechst assay was performed to compare the sensitivity between the conshRNA vs. Keap1 shRNA when cells were treated with various concentrations of either cisplatin (B) or smoke exposure (C) for 3 days. The corresponding cells were seeded at the same density on day 1 and changed to fresh media the following day (day 2). Then the conditioned media were harvested after 24, and 48 h (day 3 and day 4), and the total number of cells counted accordingly. Secreted PGE2 (D) and IL-6 levels (E) were measured using ELISA kit and normalized to cell number. All data are repeated in triplicates and represented as mean ± SD. p-values were calculated using Student's t-test. *p<0.05, **P<0.05, and ***p<0.001

#### SUPPLEMENTAL FIGURE 14

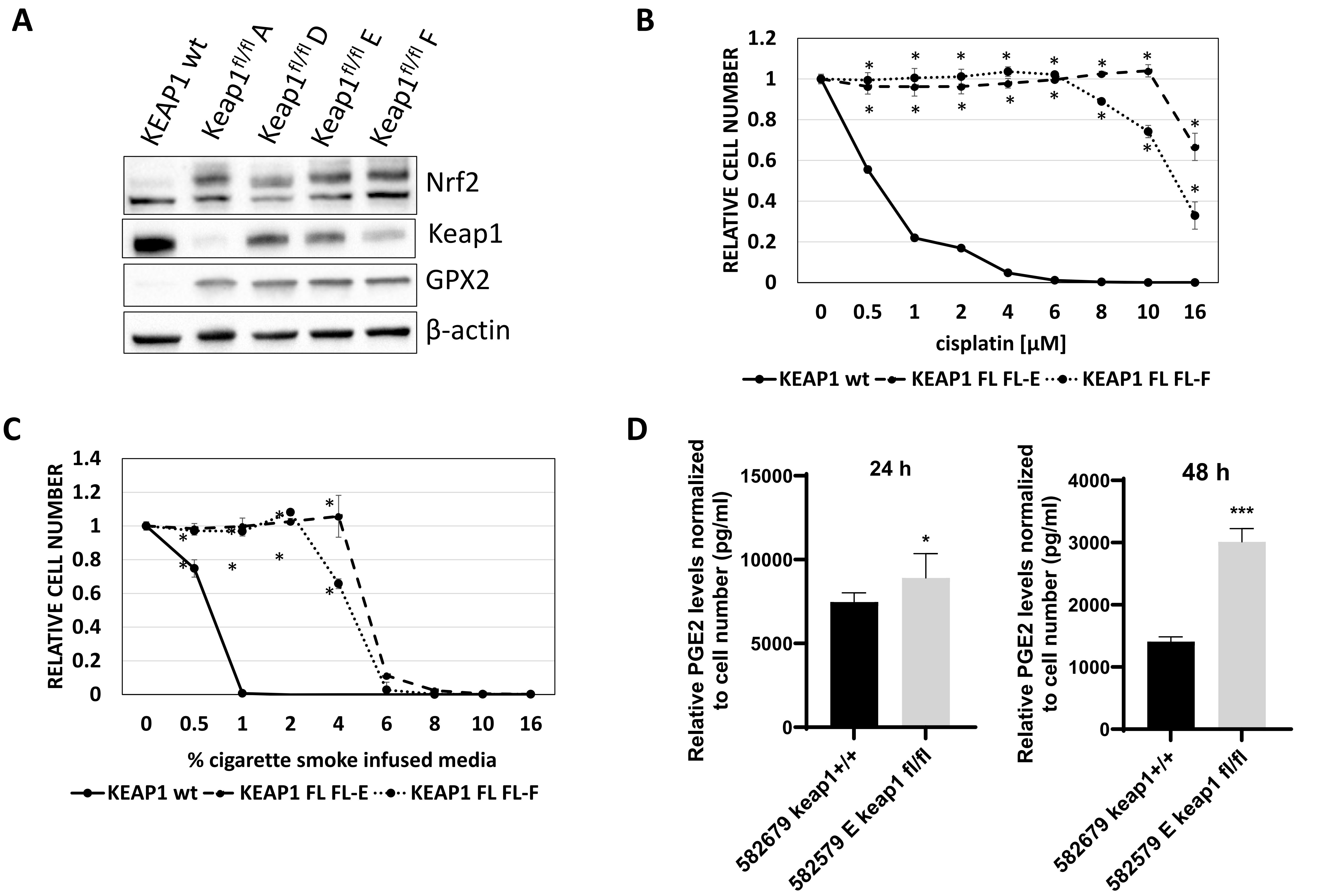


**Supplemental Figure 14. Stable Nrf2 activation through KEAP1 partial deletion.** Genetically engineered mouse models of HPV-negative HNSCC were established by frequently altered genes of TP53 and PIK3CA. Tumors were digested using a Miltenyi tumor dissociation kit, and single-cell suspension was generated. Keap1 wildtype and Keap1$^{fl/fl}$ Clones A, D, E, and F were kindly provided by Dr. Rutulkumar Patel. Keap1+/+ is a cell line with two copies of wildtype Keap1, and Keap1$^{fl/fl}$ represents one deleted and one flox copy of keap1. Clones A, D, E, and F were isolated from the Keap1$^{fl/fl}$ cells. (A) Cells were harvested for Nrf-2 and GPX2 protein expression detection. Hoechst assay was performed to compare the sensitivity between the Keap1 +/+ vs. Keap1$^{fl/fl}$ Clone E and F upon treatment with cisplatin (B) and smoke treatment (C) for 3 days. Statistically significant deviations were observed from Keap wt vs. Keap1$^{fl/fl}$ for each treatment (*p < 0.05). The corresponding cells were seeded at the same density on day 1 and changed to fresh media the following day (day 2). Then the conditioned media were harvested after 24, and 48 h (day 3 and day 4), and the total number of cells counted accordingly. Secreted PGE2 levels (D) were measured using ELISA kit and normalized to cell number. All data are repeated in triplicates and represented as mean ± SD. p-values were calculated using Student's t-test. *p<0.05, **P<0.05, and ***p<0.001

#### SUPPLEMENTAL FIGURE 15

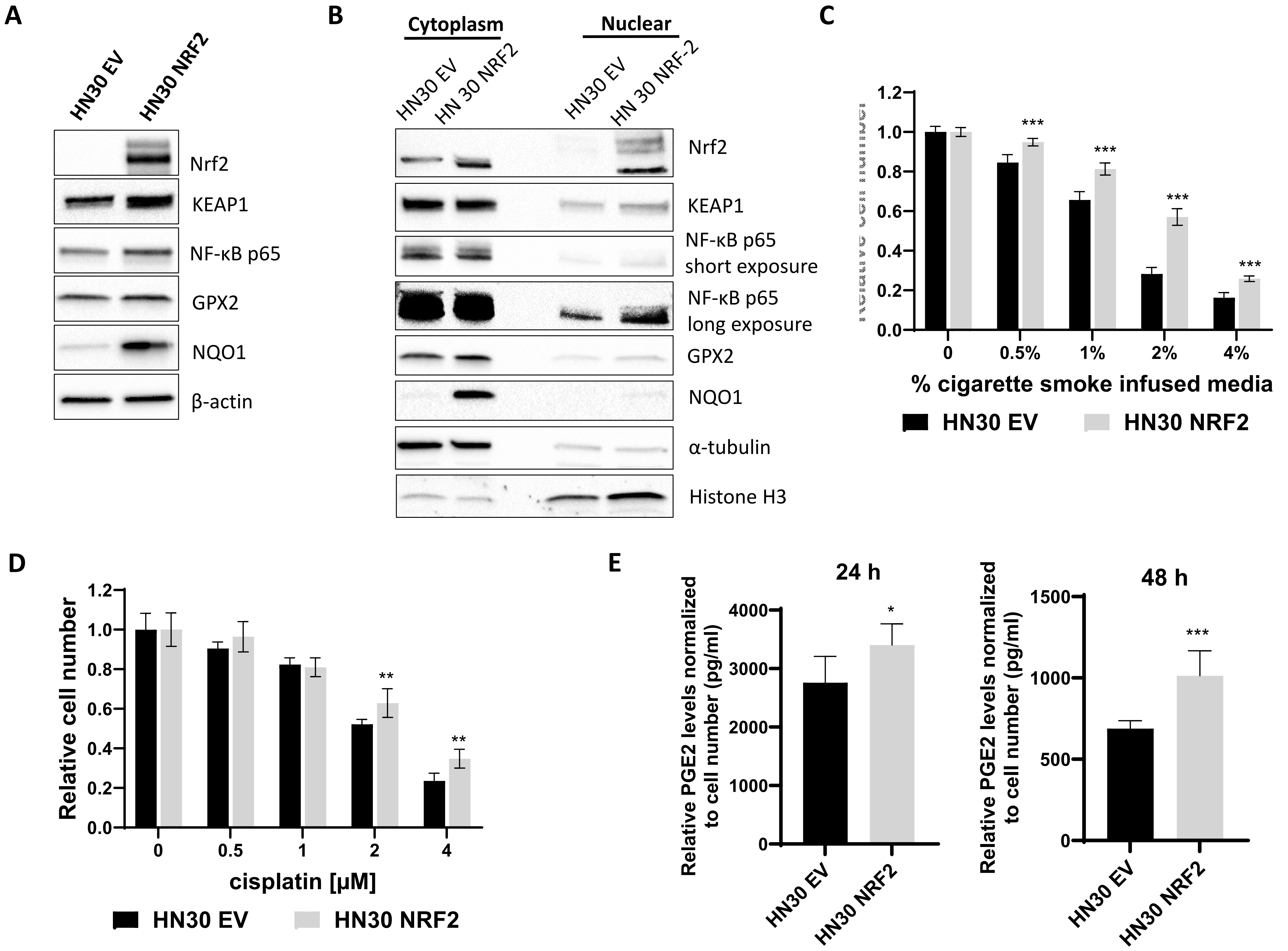


**Supplemental Figure 15. Nrf2 is linked to cisplatin resistance, NFkB activation and variable PGE2 production.** HEK293T cells were transfected with either human NRF2 overexpression (OE) plasmid or empty vector (EV) from GIPZ. The corresponding viruses were collected and infected with human HNSCC HN30 cells. After 4 µg/mL puromycin treatment for 3 days, cells were harvested for whole cell lysate, followed by Western blot analysis for the Nrf-2 targeted proteins (A). (B) Fractionation was conducted to confirm the translocation of Nrf-2 targeted proteins. α-Tubulin served as loading control for the cytoplasmic fraction, and histone H3 served as loading control for the nuclear fraction. Hoechst assay was performed to compare the sensitivity between the MOC1 EV vs. Nrf-2 OE when cells were treated with various concentrations of smoke exposure (C) and cisplatin (D) treatment for 3 days. The corresponding cells were seeded at the same density on day 1 and changed to fresh media the following day (day 2). Then the conditioned media were harvested after 24, and 48 h (day 3 and day 4), and the total number of cells counted accordingly. Secreted (E) was measured using ELISA kit and normalized to cell number. All data are repeated in triplicates and represented as mean ± SD. p-values were calculated using Student's t-test. *p<0.05, **P<0.05, and ***p<0.001

#### SUPPLEMENTAL FIGURE 16

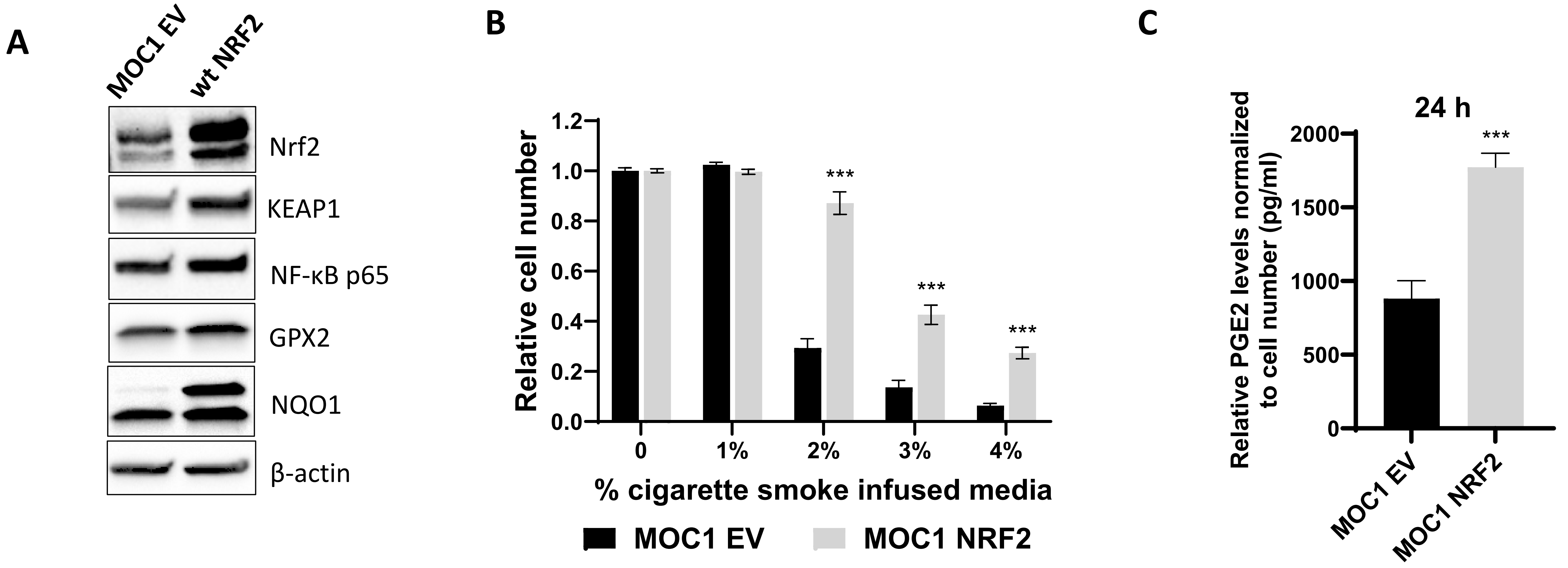


**Supplemental Figure 16. Nrf2 activation is associated with resistance to cigarette smoke exposure and altered PGE2 production.** HEK293T cells were transfected with either mouse NRF2 OE plasmid or empty vector (EV) from GIPZ. The corresponding viruses were collected and infected with HN30 (A) cells. After 4 µg/mL puromycin treatment for 3 days, cells were harvested for whole cell lysate, followed by Western blot analysis for the Nrf-2 targeted proteins. (B) Hoechst assay was performed to compare the sensitivity between the EV vs. NRF2 OE when cells were treated with various concentrations of smoke exposure for 3 days. The corresponding cells were seeded at the same density on day 1 and changed to fresh media the following day (day 2). Then the conditioned media were harvested after 24, and 48 h (day 3 and day 4), and the total number of cells counted accordingly. Secreted PGE2 (C) was measured using ELISA kit and normalized to cell number. All data are repeated in triplicates and represented as mean ± SD. p-values were calculated using Student's t-test. $*p<0.05$, $**P<0.05$, and $***p<0.001$

#### SUPPLEMENTAL FIGURE 17

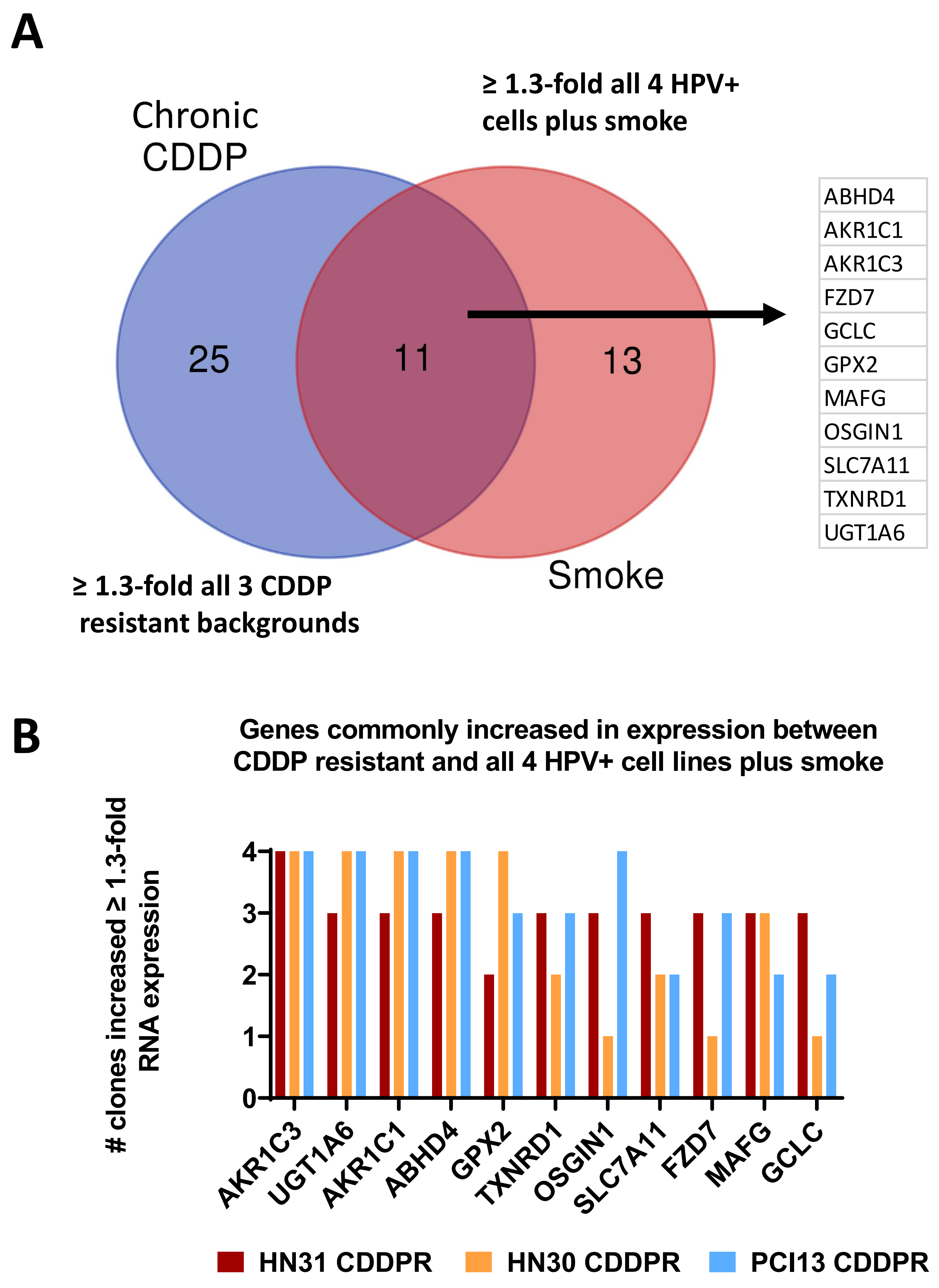


**Supplemental Figure 17. NRF2 target genes commonly upregulated in CDDP resistant and smoke exposed HNSCC cell lines.** (A) Venn diagram illustrating the number of NRF2 signature genes significantly elevated in 3 different CDDP resistant HNSCC cell line genetic backgrounds, 4 HPV+ HNSCC cell lines treated with smoke, and their overlap. A total of 36 common NRF2-target genes were significantly elevated in clones from all 3 different CDDP resistant cell lines (HN30, HN31, and PCI13) and 11 of these genes (listed to right) overlapped with all 4 HPV+ HNSCC cell lines treated with smoke. (B) Gene expression data in the above Venn diagram was derived from four different CDDP resistant clones each isolated from HN30, HN31, and PCI13 (i.e., 12 clones), and the number of clones for each cell where the 11 common NRF2 target genes was elevated is plotted in the bar graph.

### SUPPLEMENTAL FIGURE 18

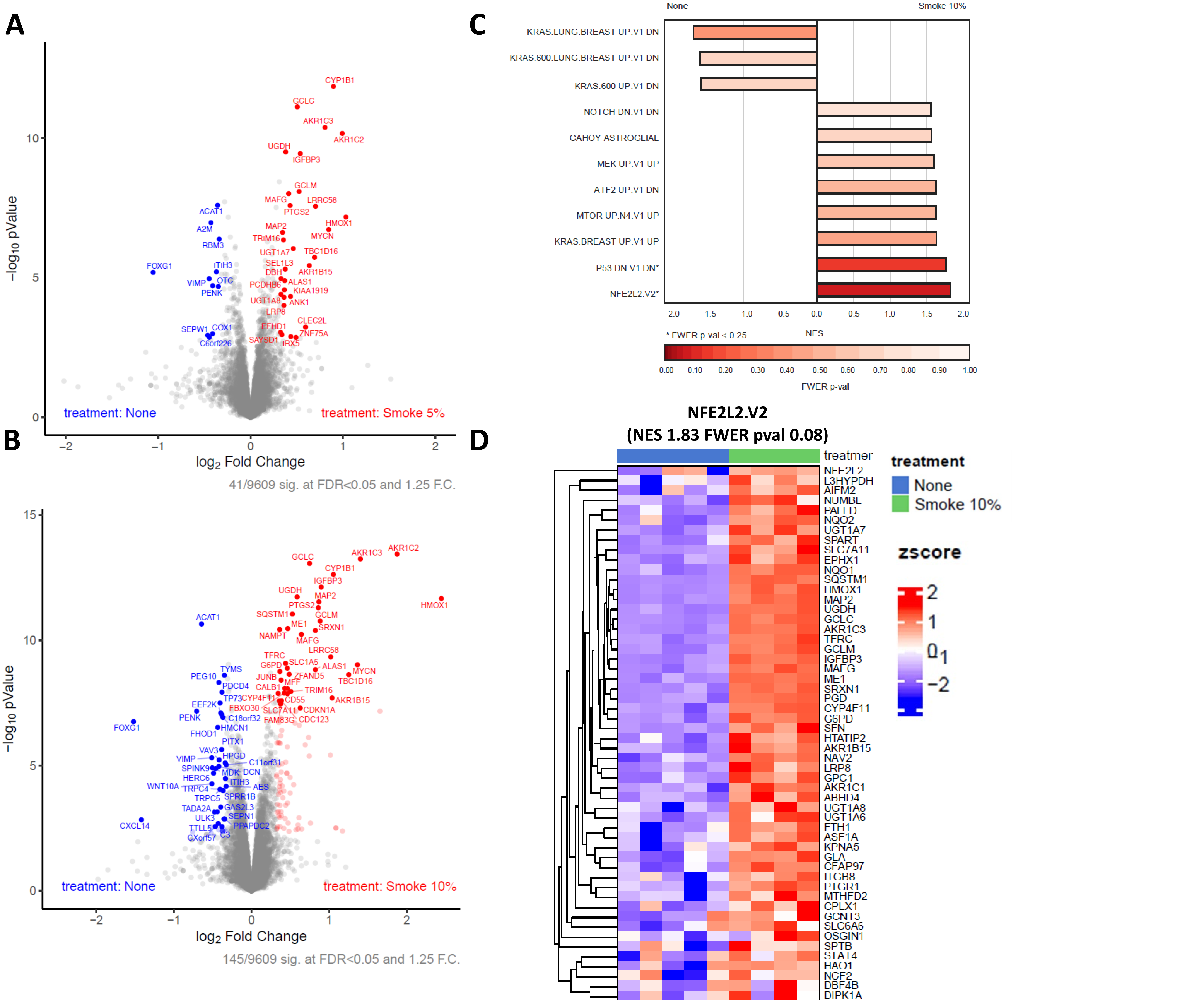


**Supplemental Figure 18. Smoke activates Nrf2 dependent protein synthesis.** UDSCC2 cells were exposed to 5% or 10% cigarette infused media for 12 hours. Cells were collected and subjected to protein analysis using mass spectrometry. Volcano plots of proteins as a function of fold change and log10 p-value at 5% (A) and 10% (B) smoke exposure. C) Pathway enrichment and depletion as a function of smoke exposure (10%). D) Nrf2 pathway protein levels (10% smoke exposure).

### SUPPLEMENTAL FIGURE 19

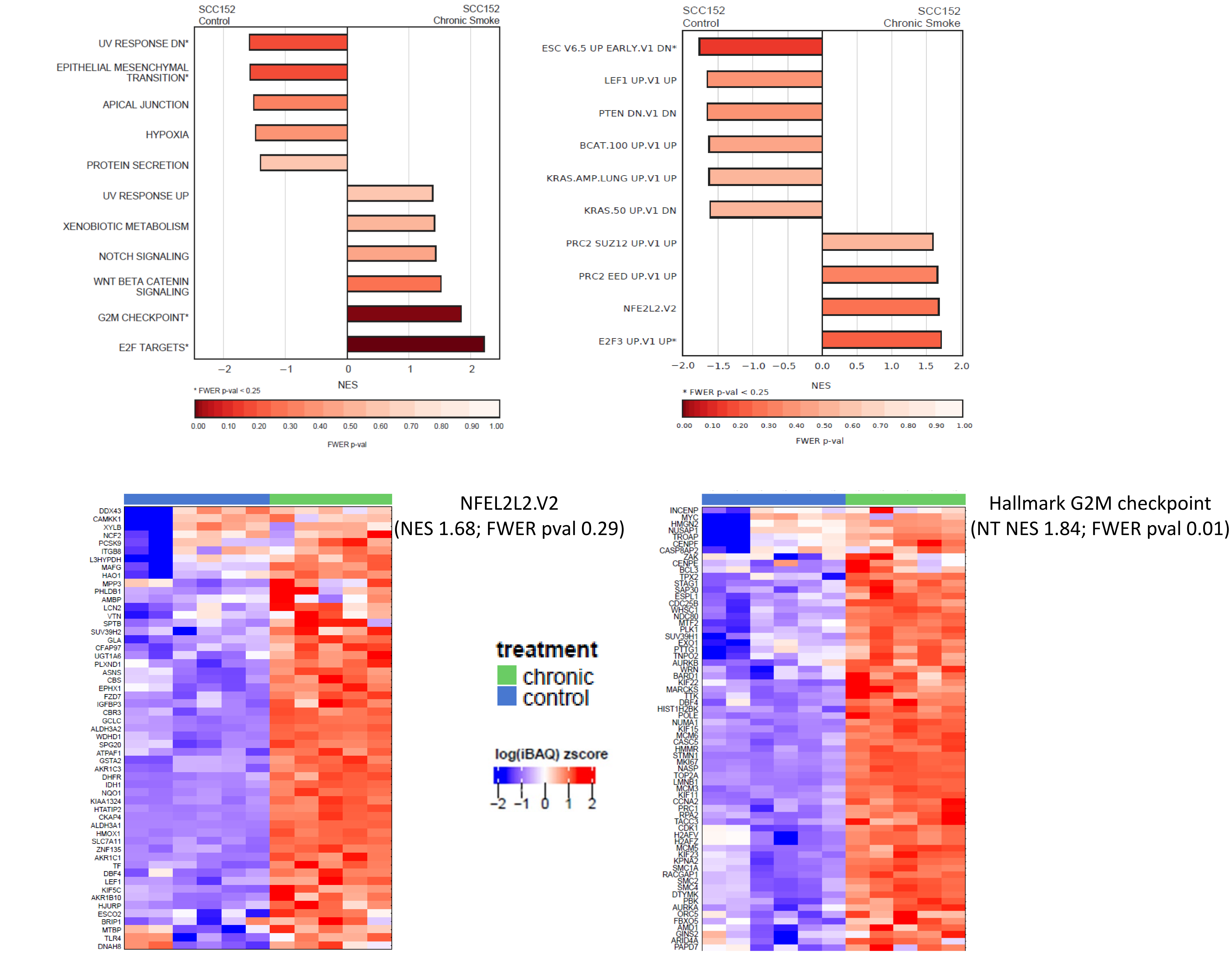


**Supplemental Figure 19. Chronic smoke exposure activates Nrf2 dependent protein synthesis and reduces proliferation.** SCC152 cells were chronically exposed to 5% cigarette smoke infused media and exposed to an acute bolus of 5% smoke infused media; proteins were isolated 12 hours following the acute exposure. Pathway enrichment analysis using the Hallmark (left) and Oncogenic 6 (right) panels. Nrf2 and G2M checkpoint pathway protein levels are illustrated below, left and right respectively.

### SUPPLEMENTAL FIGURE 20

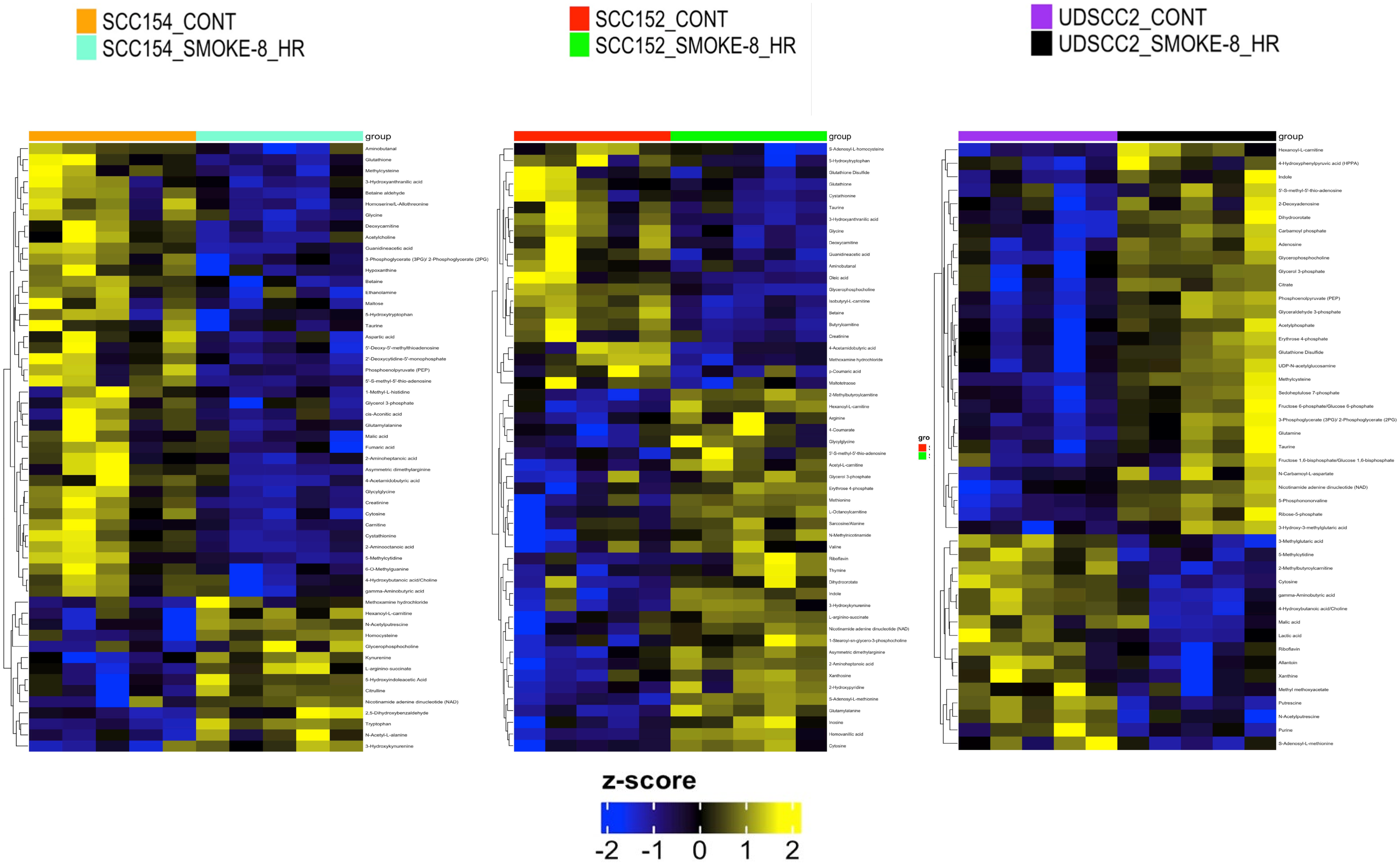


**Supplemental Figure 20. Acute smoke exposure activates metabolic shifts neutralizing oxidative stress.** Acute smoke exposure (8hr; 10%) of SCC154, SCC152 and UDSCC resulted in profound metabolic shifts, focused on pathways designed to neutralize oxidative stress and rebuild reducing equivalents.

**SUPPLEMENTAL FIGURE 21**

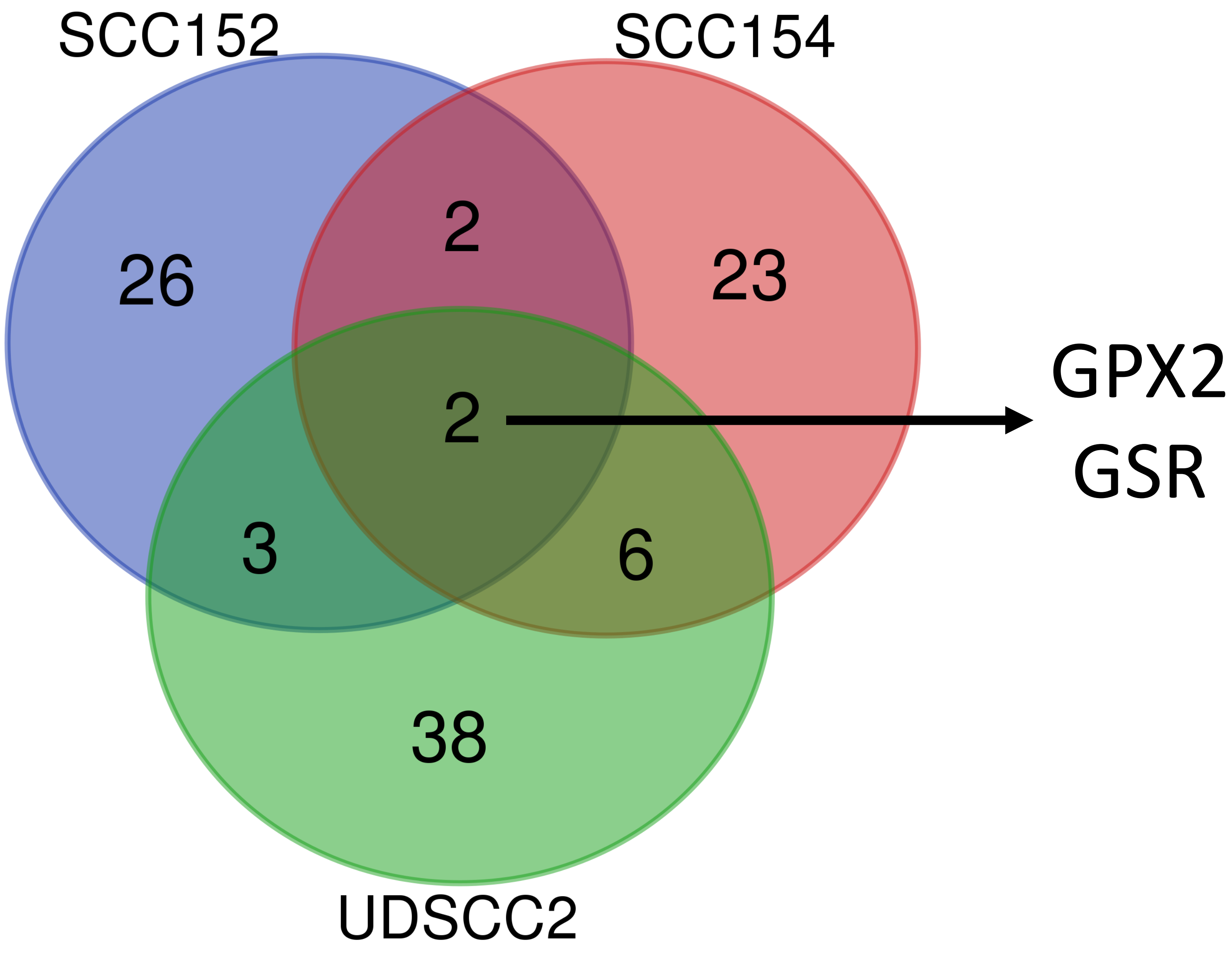


**Supplemental Figure 21. Integrated transcriptomic – metabolomic analysis of smoke exposure.** Gene expression and altered metabolite data were integrated for 3 cell lines (SCC152, SCC54, UDSCC2) using metabolic shifts identified at the 8hour time point. Genes with a FDR of ≤0.05 were compared across the 3 cell backgrounds and common genes which were altered were identified.

#### SUPPLEMENTAL FIGURE 22

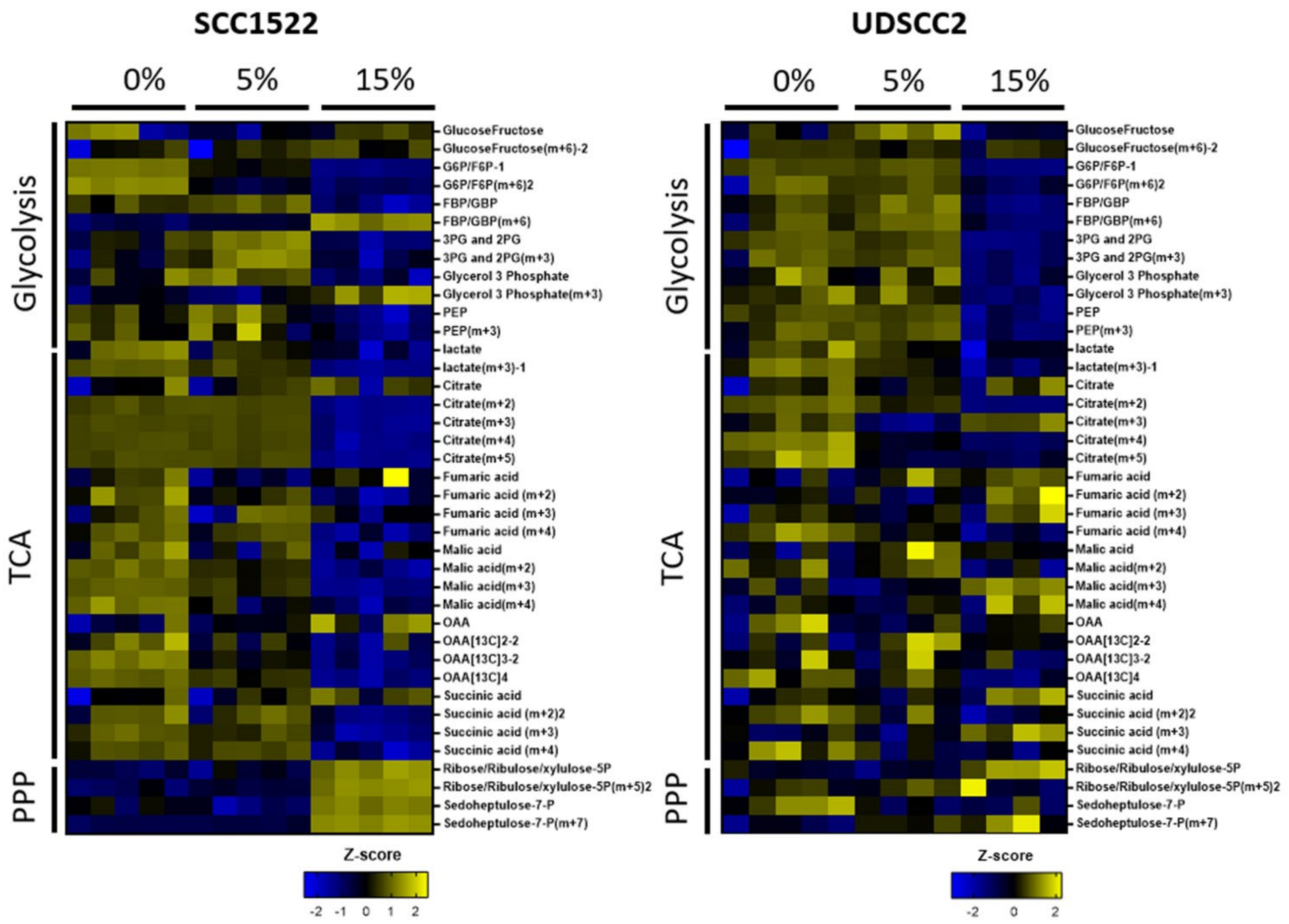


**Supplemental Figure 22. Acute smoke exposure activates metabolic shifts neutralizing oxidative stress.** SCC152 and UDSCC2 cells were exposed to smoke infused media (0%, 5%, 15%) for 8 hours in the presence of 25mM 13C all labeled glucose (Glc). Following completion of exposure, cell lysates were analyzed for 13C incorporation which predominated in pentose phosphate pathway (PPP) intermediates) in both cell lines and Krebs cycle intermediates in the case of UDSCC2.

#### SUPPLEMENTAL FIGURE 23

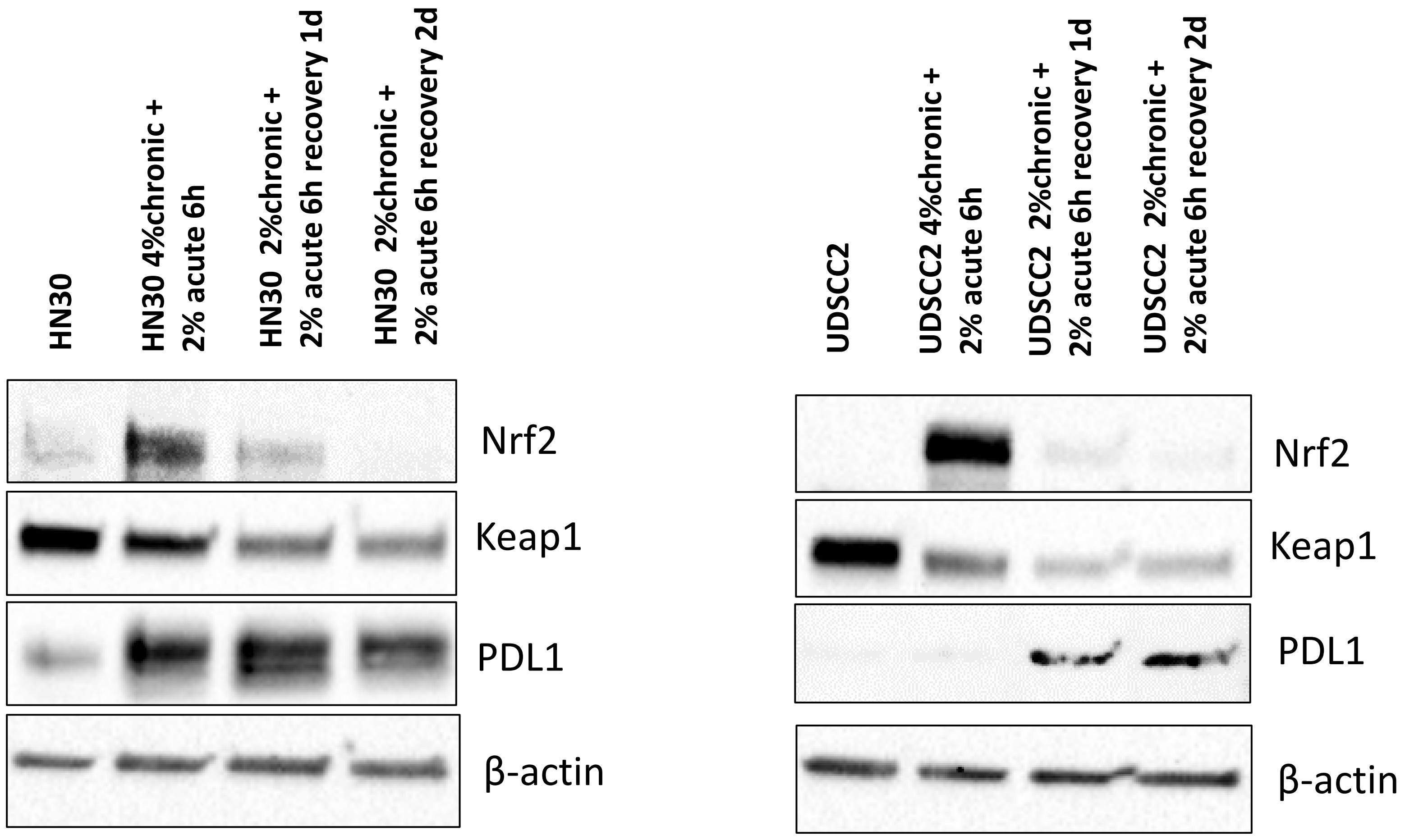


**Supplemental Figure 23. Smoke exposure, Nrf2 activation and PDL1 levels.** HPV-positive UDSCC2 and HPV-negative HN30 cells were chronically exposed to cigarette-infused media at specified concentrations for 3-6 months. An additional bolus of 4% smoke was administered for 6 hours, followed by a change to fresh media for 24 and 48 hours before harvesting for western blot analysis. β-actin was used as the protein loading control.

### SUPPLEMENTAL FIGURE 24

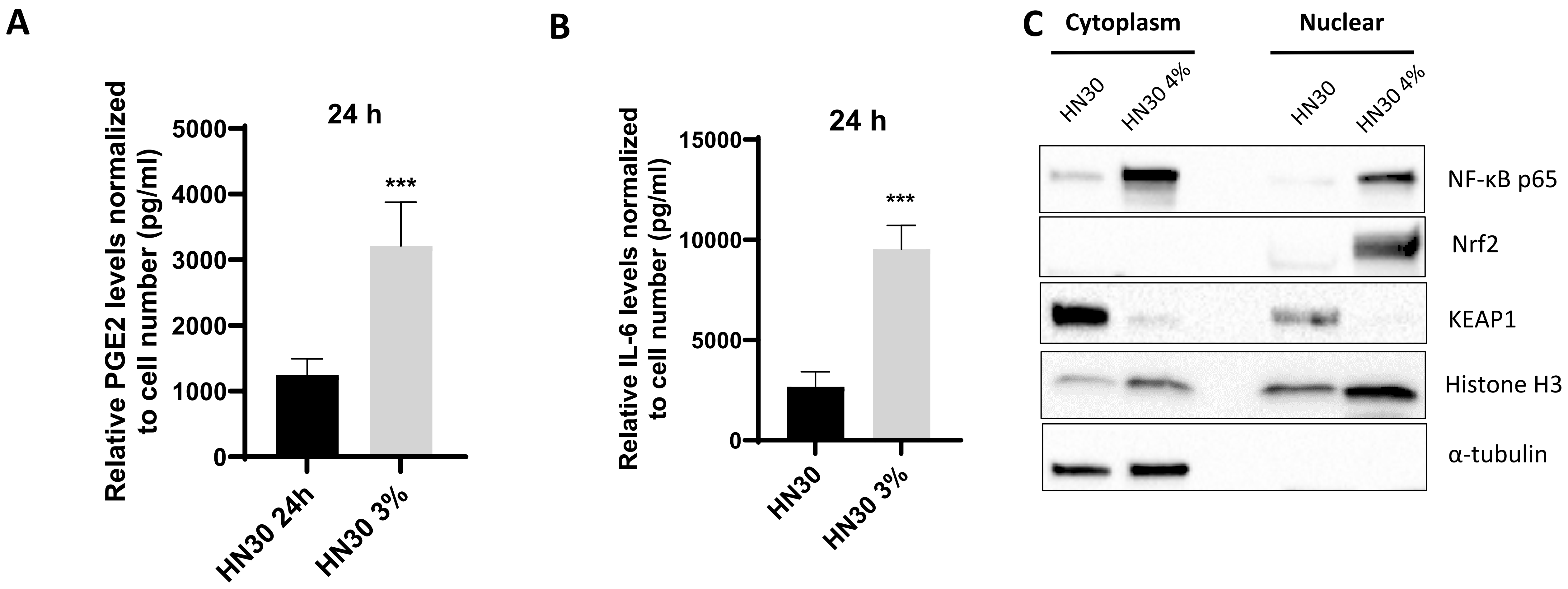


**Supplemental Figure 24. Chronic smoke exposure and immune modulation.** The corresponding cells were seeded at the same density on day 1 and changed to fresh media the following day (day 2). Then the conditioned media were harvested after 24, and 48 h (day 3 and day 4), and the total number of cells counted accordingly. Secreted PGE2 (A) and IL-6 levels (B) were measured using ELISA kit and normalized to cell number. All data are repeated in triplicates and represented as mean ± SD. p-values were calculated using Student's t-test. *p<0.05, **P<0.05, and ***p<0.001

### SUPPLEMENTAL FIGURE 25

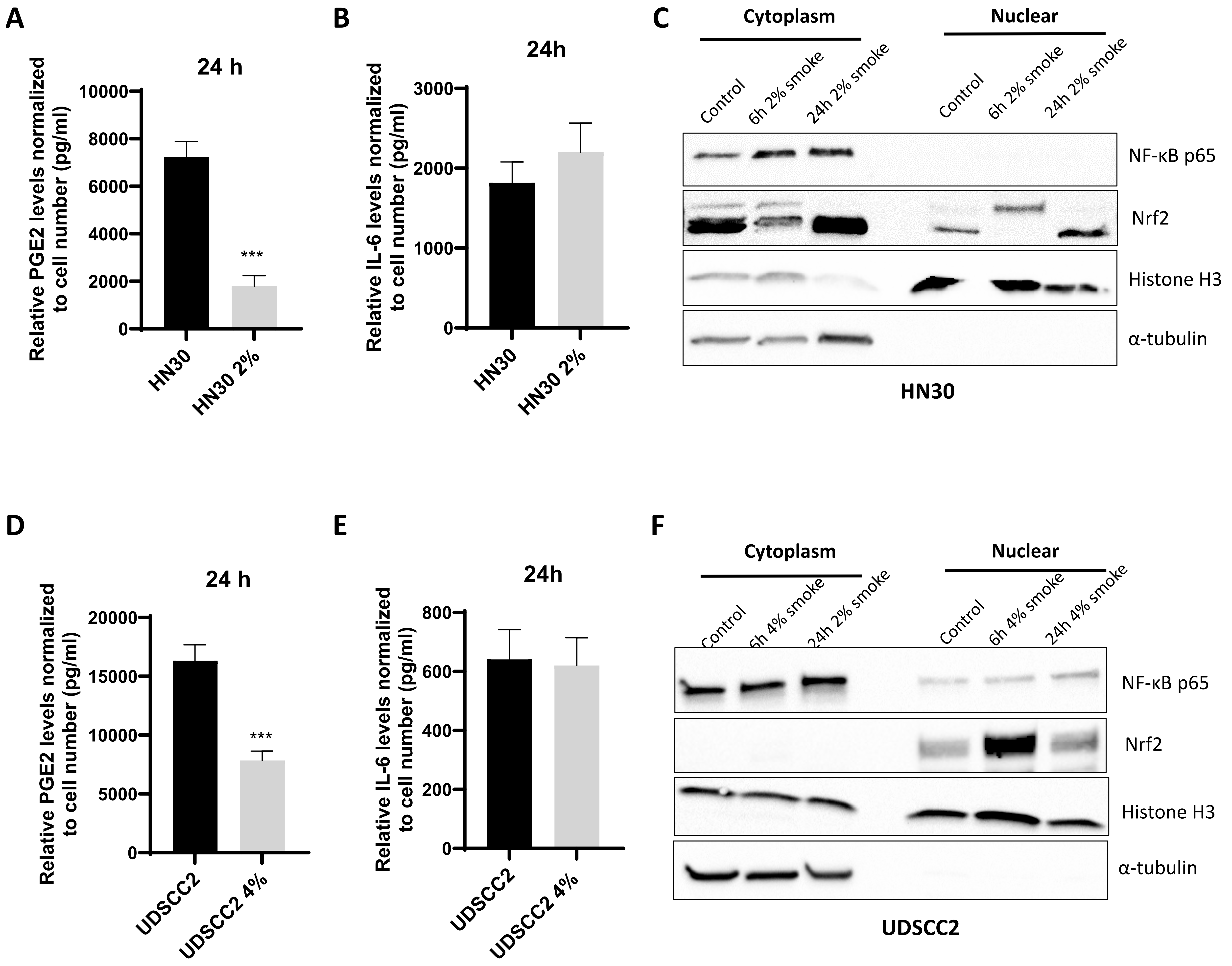


**Supplemental Figure 25. Acute smoke exposure and immune modulation.** The corresponding cells were seeded at the same density on day 1 and treated with or without 2% smoke media the following day (day 2). Then the conditioned media were harvested after 24 h (day 3), and the total number of cells counted accordingly. Secreted PGE2 (A, D) and IL-6 (B, D) levels were measured using the corresponding ELISA kit and normalized to cell number. All data are repeated in triplicates and represented as mean ± SD. p-values were calculated using Student's t-test. (*P<0.05, **P<0.05, and ***p<0.001). (C, F) Fractionation was conducted to confirm the translocation of Nrf-2 targeted proteins. α-Tubulin served as loading control for the cytoplasmic fraction, and histone H3 served as loading control for the nuclear fraction. *p<0.05, **P<0.05, and ***p<0.001

#### SUPPLEMENTAL FIGURE 26

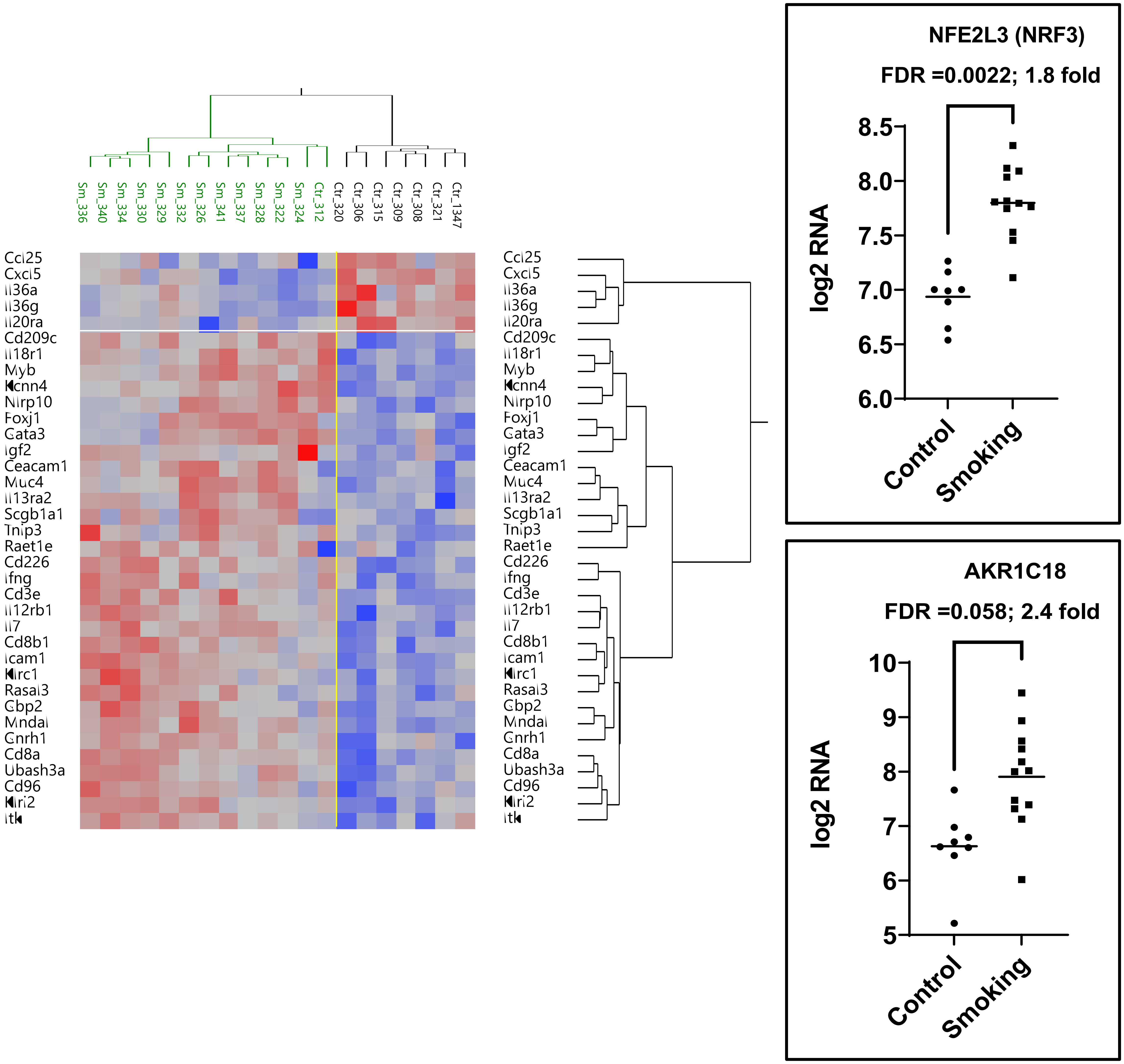


**Supplemental Figure 26. Chronic smoke exposure and immune modulation.** MOC1 cells were used to generate orthotopic tongue xenografts in immunocompetent mice. MOC1 cells chronically exposed to 4% smoke were used to generate orthotopic tongue xenografts in chronically smoke exposed immunocompetent mice. Tumors were harvested, snap frozen and subjected to RNAseq analysis.

#### SUPPLEMENTAL FIGURE 27

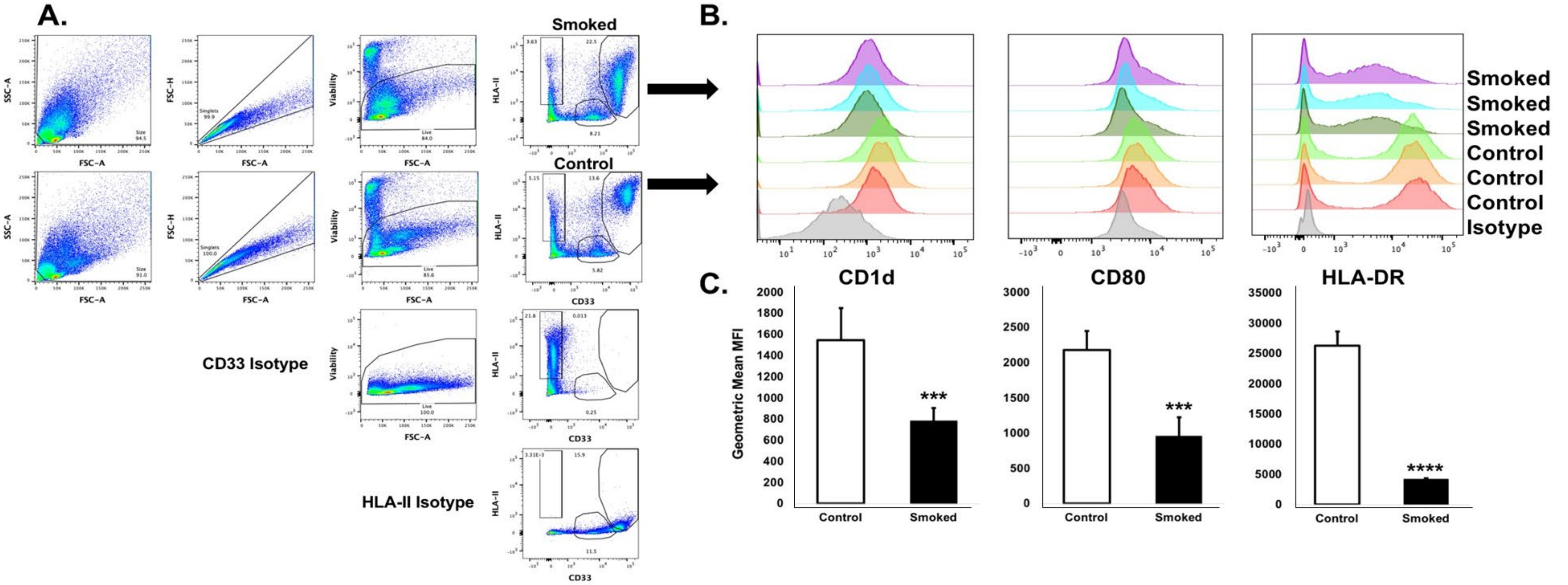


**Supplemental Figure 27. Smoked –exposed tumor cell culture media inhibits expression of myeloid cell antigen presentation mediators.** HN30 tumor cells exposed to tobacco smoke chronically (4%) followed by an acute bolus (4%) and HN30 cells exposed to filtered air were used to generate conditioned media for 24 hours. Cell culture media was harvested and added to PBMC derived from three healthy donors which were cultured for an additional 48 hours. Cells were then harvested and characterized by flow cytometry. **A)** Myeloid cell gating strategy. **B)** Flow cytometry histograms of antigen presentation marker expression CD1d, CD80, and HLA-DR. **C)** Graphical quantitation of flow histograms shown in **B.** ***$p<0.005$, ****$p<0.001$ by Student's paired two-tailed t-test.
